# The mouse cortico-basal ganglia-thalamic network

**DOI:** 10.1101/2020.10.06.326876

**Authors:** Nicholas N. Foster, Laura Korobkova, Luis Garcia, Lei Gao, Marlene Becerra, Yasmine Sherafat, Bo Peng, Xiangning Li, Jun-Hyeok Choi, Lin Gou, Brian Zingg, Sana Azam, Darrick Lo, Neda Khanjani, Bin Zhang, Jim Stanis, Ian Bowman, Kaelan Cotter, Chunru Cao, Seita Yamashita, Amanda Tugangui, Anan Li, Tao Jiang, Xueyan Jia, Zhao Feng, Sarvia Aquino, Gordon Dan, Marina Fayzullina, Hyun-Seung Mun, Sarah Ustrell, Tyler Boesen, Anthony Santarelli, Muye Zhu, Nora L. Benavidez, Monica Song, David L. Johnson, Hanpeng Xu, Michael S. Bienkowski, X. William Yang, Hui Gong, Ian Wickersham, Qingming Luo, Byung Kook Lim, Li I. Zhang, Houri Hintiryan, Hongwei Dong

## Abstract

The cortico-basal ganglia-thalamic loop is one of the fundamental network motifs in the brain. Revealing its structural and functional organization is critical to understanding cognition, sensorimotor behavior, and the natural history of many neurological and neuropsychiatric diseases. Classically, the basal ganglia is conceptualized to contain three primary information output channels: motor, limbic, and associative. However, given the roughly 65 cortical areas and two dozen thalamic nuclei that feed into the dorsal striatum, a three-channel view is overly simplistic for explaining the myriad functions of the basal ganglia. Recent works from our lab and others have subdivided the dorsal striatum into numerous functional domains based on convergent and divergent inputs from the cortex and thalamus. To complete this work, we generated a comprehensive data pool of ∼700 injections placed across the striatum, external globus pallidus (GPe), substantia nigra pars reticulata (SNr), thalamic nuclei, and cortex. We identify 14 domains of SNr, 36 in the GPe, and 6 in the parafascicular and ventromedial thalamic nuclei. Subsequently, we identify 6 parallel cortico-basal ganglia-thalamic subnetworks that sequentially transduce specific subsets of cortical information with complex patterns of convergence and divergence through every elemental node of the entire cortico-basal ganglia loop. These experiments reveal multiple important novel features of the cortico-basal ganglia network motif. The prototypical sub-network structure is characterized by a highly interconnected nature, with cortical information processing through one or more striatal nodes, which send a convergent output to the SNr and a more parallelized output to the GPe; the GPe output then converges with the SNr. A domain of the thalamus receives the nigral output, and is interconnected with both the striatal domains and the cortical areas that filter into its nigral input source. This study provides conceptual advancement of our understanding of the structural and functional organization of the classic cortico-basal ganglia network.

## INTRODUCTION

The basal ganglia is a collection of interconnected cerebral nuclei that receive major projections from all parts of the cortex (Oh et al. 2014, Hintiryan, Foster et al. 2016; Hunnicutt et al. 2016), making it the chief subcortical array for processing cortical information. The principal components of the basal ganglia are the striatum, pallidum, and substantia nigra, and they have been associated with functions as diverse as ramp movements (DeLong & Strick, 1974), reinforcement learning (Balleine & O’Doherty, 2010), executive function (Dirnberger & Jahanshahi, 2013; Harrington et al. 2014), visual processing (Ding & Gold, 2013), and perception of passage of time (Lusk & Buonomano, 2016; Bakhurin et al. 2017). Aberrant basal ganglia functions are implicated in movement disorders (Gittis & Kreitzer, 2012), neuropsychiatric disorders (Graybiel & Rauch, 2000; Levy & Dubois, 2006; Gunaydin & Kreitzer, 2016; Haroon et al. 2016; Vaghi et al. 2017), and drug addiction (Koob & Volkow, 2016; Everitt & Robbins, 2016). Now-classic works have revealed that the basal ganglia is one stage in a serial circuit from cortex to basal ganglia to thalamus and back up to cortex, or the cortico-basal ganglia-thalamo-cortical loop (Nauta & Mehler, 1961; Szabo, 1962; Gerfen, 1984, 1985; Deniau & Chevalier, 1985; Alexander, et al. 1986; Gimenez-Amaya & Graybiel, 1990; Hedreen & DeLong, 1991; Hoover & Strick, 1993; Parent & Hazrati, 1994). This recurrent network is one of the most important structural motifs in the mammalian brain, with nearly half of the brain participating in its elements.

Identification of the specific subnetworks within this archetypal model is crucial to understanding how the multitude of functions and pathologies ascribed to the basal ganglia is governed. The consensus view that has emerged from both connectional and functional studies is that there are three parallel channels of information flow through the basal ganglia: associative, limbic, and sensorimotor (Alexander et al. 1986; Parent & Hazrati, 1995; Haber, 2003; Wallace et al. 2017; Mandelbaum et al. 2019; Aoki et al. 2019) in roughly the dorsomedial, ventromedial, and lateral two quadrants, respectively, of the commissural level of the caudoputamen (CP; Figure 1A). We recently described a multi-scale network organization of the mouse corticostriatal projectome (Hintiryan, Foster et al. 2016), which partitioned the territory of the CP into fine-scale subnetworks (termed *domains*) nested within intermediate-scale networks (termed *communities*) nested within large-scale networks (termed *divisions*) (Figure 1A,B). Our community-scale network appears to correspond well with the quadrant layout of the three major channels in the CP. However, each community comprises domains that each receive a unique set of cortical inputs, a finding that strongly suggests there must be a more refined, granular level of organization than the classic associative, limbic, and sensorimotor networks (Figure 1C). Previous work has attempted to identify more specific subnetworks. A meta-analytic-style approach combining results of transsynaptic retrograde tracing, classical neuronal tracing, and electrophysiology experiments obtained in primate models has been used to postulate a number of specific parallel cortico-basal ganglia subnetworks or “output channels” (Alexander et al. 1986; Alexander et al. 1990; Hoover & Strick, 1993; Middleton & Strick, 2001). However, accurate assessment of the number of pathways has been limited by an inadequate knowledge of the organization of the striatum, which has at best been alternately conceptualized according to the caudate/putamen divisions or the quadrant schema.

**Figure 1.**
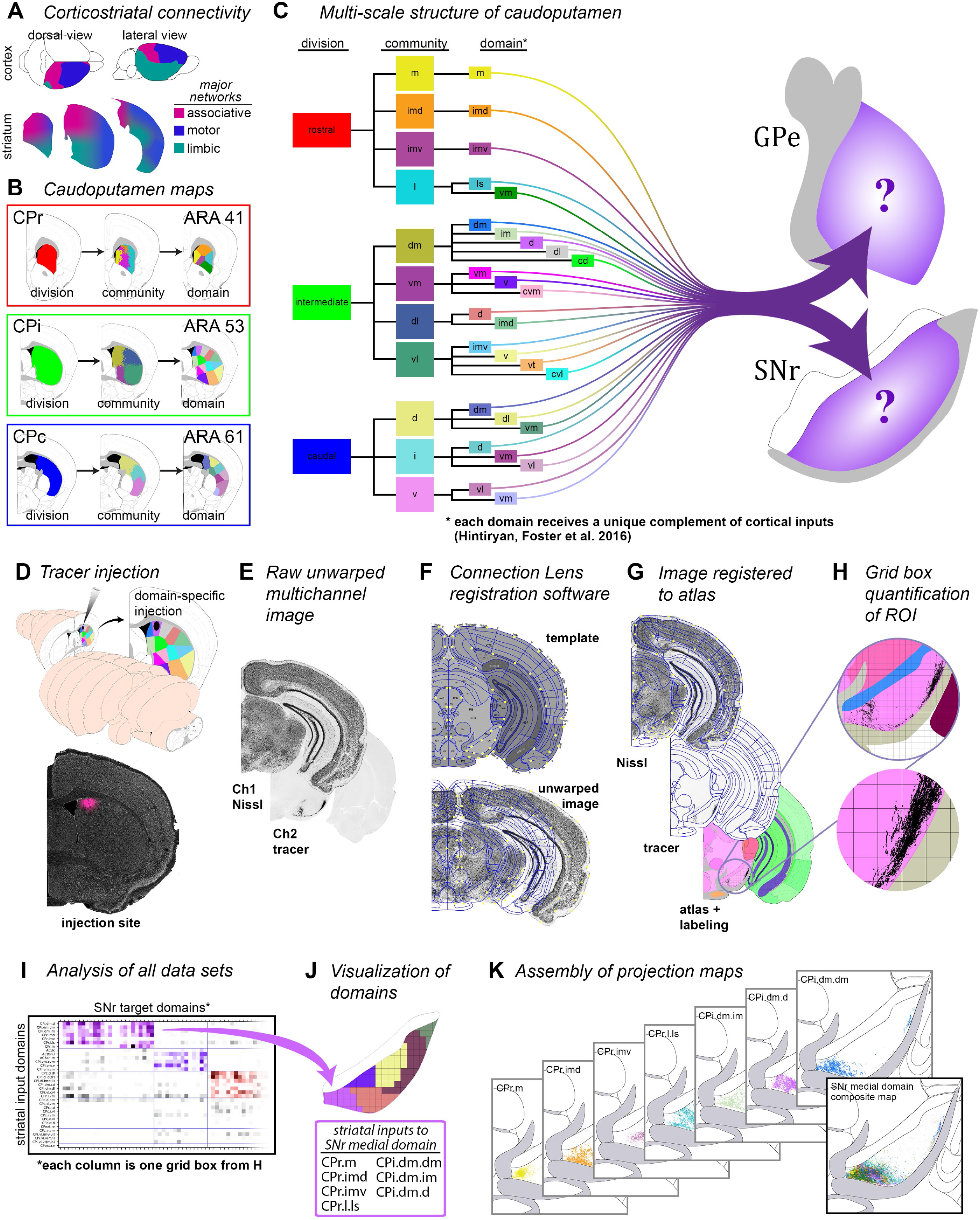
Overview and workflow. A) General topography of the 3 classic corticostriatal macrocircuits: motor, limbic, and associative. B) Map of the multi-scale subdivisions of the caudoputamen at rostral (CPr), intermediate (CPi), and caudal (CPc) levels. C) Dendrogram of the multi-scale, hierarchical structure of the CP, depicting how each division is composed of smaller communities and even smaller domains, each with a unique set of cortical inputs. The topography of the striatofugal pathway is unknown. The data production workflow starts with D) discrete injection of anterograde tracer into one of the striatal domains. Tissue sections are E) imaged and imported into F) Connection Lens where fiducial points in the nissl channel are matched to the atlas template; G) images are deformably warped and the tracer channel is segmented into a binary threshold image. H) The software subdivides all brain regions into a square grid space and quantifies the pixels of axon labeling in each grid box. I) The quantified axonal terminals from all injections to all grid boxes at each nucleus-level is visually summarized in a matrix (darker shading indicates denser termination; colors correspond to particular domains). Statistical analysis reveals the subnetworks, groups of striatal domains that project to a common set of grid boxes. Those grid boxes in the SNr maps are then (J) colored to visualize the new input-defined pallidal or nigral domain. (K) Composite projection maps of the colored axonal terminals illustrate the striatofugal terminal pattern, with color matching the striatal source.

Our description of the domain-level organization of the corticostriatal projectome presents an entry point to parsing the entire subnetwork organization of the mouse cortico-basal ganglia loop. By following the efferent pathways of the CP domains, we identify the major output channels of the mouse globus pallidus and substantia nigra, finding 14 domains through the substantia nigra pars reticulata (SNr) and 36 domains through the globus pallidus external (GPe). Further following the efferents of some of the domains of the SNr through the thalamus reveals 6 output channels traversing the parafascicular nucleus (PF) and 6 through the ventromedial nucleus (VM) with different thalamocortical innervation patterns between the two nuclei. We characterize the dendritic morphology of SNr neurons, as a further index of how striatal inputs may be converging in the nigra. Using the double co-injection technique, we for the first time definitively demonstrate entire continuous loops (the “loop” structure) of two of the anatomical subnetworks, which until now had only been inferred by combining experimental results demonstrating segments of loops. Using channelrhodopsin-assisted circuit mapping, we demonstrate for the first time that the thalamic feedback up to the cortex synapses directly onto the corticostriatal neurons that feed into that subnetwork loop, forming a true closed-circuit loop. We also demonstrate a direct corticonigral pathway using Cre-dependent neuronal tracing and specific visualization of corticonigral terminal boutons. Collectively, these various approaches make clear the exquisite specificity and interconnectedness within the cortico-basal ganglia-thalamo-cortical subnetworks, and provide conceptual advancement for us to understand the structural and functional organization of the entire central nervous system.

## RESULTS

Injections of anterograde tracers were made into each of the domains of the CP (Hintiryan, Foster et al. 2016) as well as the core, medial, and lateral shell of the nucleus accumbens, for a total of 33 injections (Figure 1D). Coronal sections containing the SNr and GPe were photographed and images were registered to a common anatomic atlas, the Allen Reference Atlas (Dong, 2007), using our in-house registration software Connection Lens (Hintiryan, Foster et al. 2016). The axonal labeling was segmented and digitally reconstructed into a binary threshold image (Figure 1E-G). The axonal reconstructions falling within the basal ganglia nuclei were quantified using a quadrat method, wherein the area of the SNr and GPe were subdivided into a square grid space, and the axons falling within each grid box were quantified (Figure 1H). This permitted an analysis not just of the density of axonal terminations in each target nucleus, but also of the subregional localization of those terminations within each nucleus. These data were then subjected to network structure analysis using the Louvain community detection algorithm (see Methods for a detailed description), which partitioned the CP domains and their terminal zones into communities, i.e., subnetworks or output channels. CP domains that projected to the same set of boxes were grouped together as a subnetwork (Figure 1I), and the set of boxes comprising the terminal zone for that subnetwork were demarcated as a new domain of the nigra or pallidum (Figure 1J). Finally, the thresholded axonal reconstruction images were re-colored with unique colors corresponding to their striatal domains of origin and collectively projected onto atlas maps to make projection maps that demonstrate the underlying axonal data that make up each new domain (Figure 1K). All of these procedures are described in more detail in Methods.

### Complex convergence and divergence of the striatonigral pathway

Striatonigral projections from each CP domain and accumbens terminate in discrete terminal zones within the SNr that are restricted mediolaterally and dorsoventrally, but most (75%) span the entire rostrocaudal extent of the SNr, forming longitudinal columns of terminal axons (Figure 2B,G,H; Supplementary Figure 1A,B; Table 1). The gross topographical organization of the striatonigral pathway is that the efferent projections along the rostrocaudal (CPr-CPi-CPc) axis of the striatum project to the medial-central-lateral SNr, respectively (Figure 2A,C, Supplementary Figure 2A), while the dorsoventral axis of the striatum is inverted in its nigral terminations (Figure 2C). These general topographic trends align well with those seen in rat and monkey (Gerfen, 1985; Hedreen & DeLong, 1991; Parent & Hazrati, 1994). Axons from CPr converge with axons from the other striatal divisions along the whole extent of the medial SNr: some CPi axons project medially and converge with CPr axons at levels 81-87, and both CPi and CPc send some axons to medial SNr at levels 89 and 91 (Figure 2A; Supplementary Figure 2A). Efferents from the CPc remain mostly segregated from the other striatal efferents, converging primarily with axons from the caudal extreme (CPext, also known as the tail) (Supplementary Figure 2C). At the community level within each striatal division, the mediolateral axis is maintained, with medial CP projecting to more medial SNr, and lateral CP to more lateral SNr, while the dorsoventral axis is inverted, with dorsal CP projecting to ventral SNr (Figure 2C,D). Consequently, we observe that axons arising from the sensory associative and limbic regions of the CPi (the communities CPi.dorsomedial (CPi.dm) and CPi.ventromedial (CPi.vm), respectively) are intermingled with axons arising from the CPr within the medial domains of the SNr, while the axons arising from the sensorimotor regions of the CPi (CPi.dorsolateral (CPi.dl) and CPi.ventrolateral CPi.vl)) remain more segregated in the middle of the SNr, forming a somatotopic map there (Figure 2F). Triple retrograde injections in the SNr precisely back-label the three communities of the CPc, confirming the specificity and discreteness of the community-level striatonigral projections (Figure 2D). Importantly, those 3 communities of the CPc were originally defined by cortical inputs (Hintiryan, Foster et al. 2016), while the labeling in Figure 2D is produced by striatal output patterns, affirming the community structure we previously described and extending that higher-order organizational scheme further through the basal ganglia.

**Figure 2.**
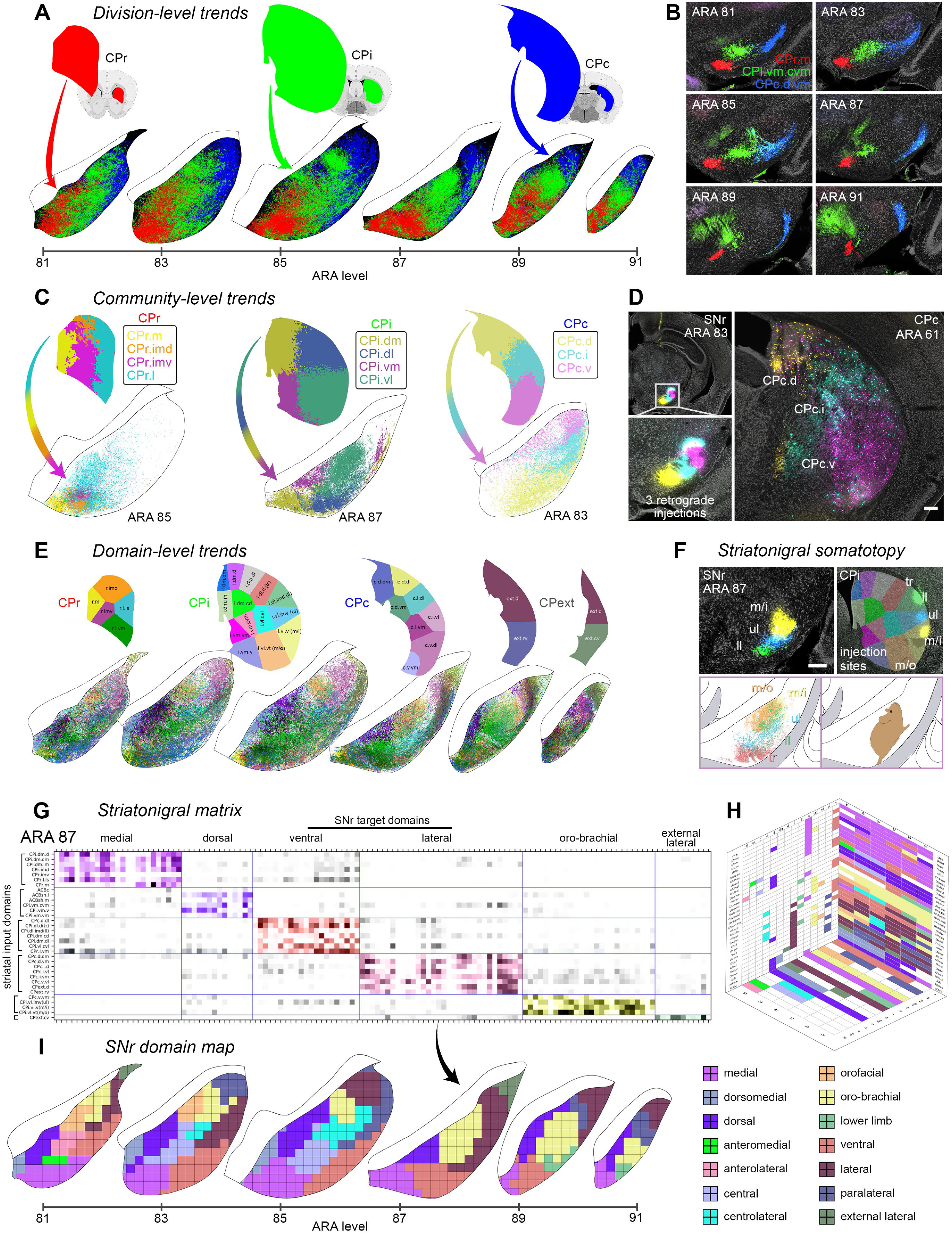
Projection trends of the striatonigral pathway at different scales and novel SNr domain map. A) Axonal reconstructions originating from all striatal domains of each CP division are colored red, green, or blue, accordingly. The rostrocaudal axis of the CP terminates in a mediolateral pattern in the SNr, respectively. B) Triple anterograde injection experiment shows labeling from rostral, intermediate, and caudal CP in a single brain. C) Community-level view of striatal terminations reveals that the dorsoventral axis of the CP terminates in an inverted pattern in the SNr. D) Triple retrograde injection experiment precisely back-labels the 3 communities of the CPc. E) Domain-level striatonigral terminations are colored according to their CP source domain. F) Triple anterograde experiment shows the lower limb (ll), upper limb (ul), and inner mouth (m/i) terminal fields in the SNr, as well as a composite reconstruction map of the full somatotopy (with trunk (tr) and outer mouth (m/o) regions as well). G) The striatonigral matrix for ARA 87 shows the density of terminations from each striatal input domain to every grid box in SNr at Atlas Level 87 (arrow points to its map in I). H) A 3D matrix represents the relationships between striatal source domains, SNr target domains, and Atlas Level of the SNr (see Supplementary Figure 1B). I) The novel SNr domain map, defined by striatonigral input topography (see Supplementary Figure 3). Scalebars = 200 µm.

**Table 1.**
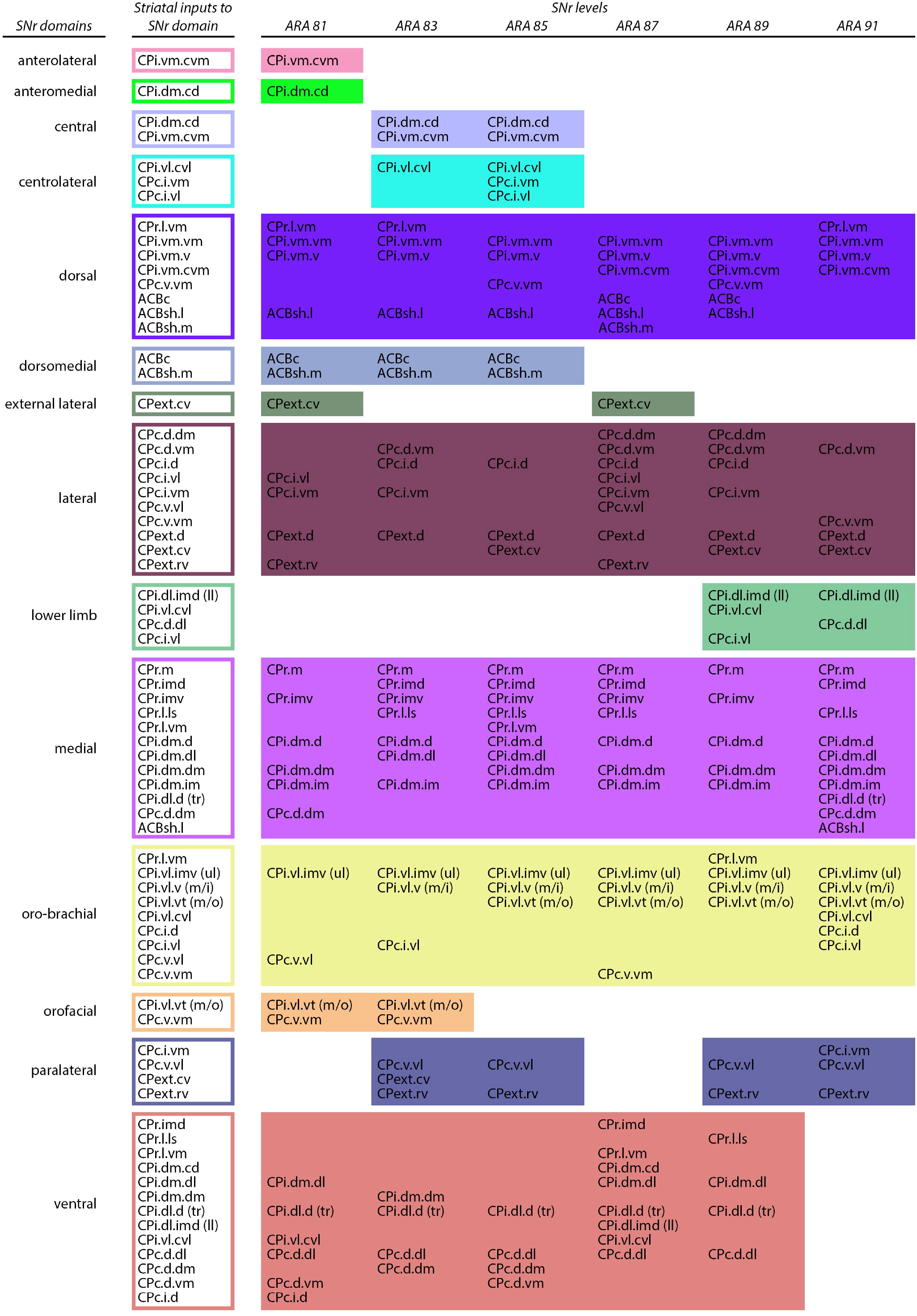
Striatal inputs to SNr domains by level

**Table 2.** Striatal inputs to GPe domains by level

| GPe domains | Striatal inputs to GPe domain | GPe levels |  |  |  |  |  |
| --- | --- | --- | --- | --- | --- | --- | --- |
|  |  | ARA 58 | ARA 60 | ARA 62 | ARA 64 | ARA 66 | ARA 68 |
| 1 | CPI.vm.vm<br>CPc.i.vm<br>ACBc<br>ACBsh.m | CPI.vm.vm<br>ACBc<br>ACBsh.m | CPI.vm.vm<br>CPc.i.vm<br>ACBc |  |  |  |  |
| 2 | CPr.imv | CPr.imv |  |  |  |  |  |
| 3 | CPr.m<br>CPI.dm.d<br>CPI.dm.dm<br>CPI.dm.im<br>CPc.d.dm | CPr.m<br>CPI.dm.dm<br>CPI.dm.im | CPr.m<br>CPI.dm.d<br>CPI.dm.dm | CPI.dm.d<br>CPI.dm.dm<br>CPI.dm.im<br>CPc.d.dm |  |  |  |
| 4 | CPr.imd<br>CPI.dm.d | CPr.imd<br>CPI.dm.d |  |  |  |  |  |
| 5 | CPI.dm.cd<br>CPI.vm.cvm<br>CPc.d.dm | CPI.dm.cd<br>CPI.vm.cvm<br>CPc.d.dm | CPI.vm.cvm<br>CPc.d.dm |  |  |  |  |
| 6 | CPr.imd<br>CPI.dl.d (tr)<br>CPI.dl.d.r<br>CPI.dm.dl<br>CPI.vl.cvl<br>CPc.d.dl | CPI.dl.d.r<br>CPI.dm.dl<br>CPI.vl.cvl | CPr.imd<br>CPI.dl.d (tr)<br>CPI.dl.d.r<br>CPI.dm.dl<br>CPc.d.dl |  |  |  |  |
| 7 | CPr.l.ls | CPr.l.ls |  |  |  |  |  |
| 8 | CPI.vl.imv.r<br>CPc.i.vm | CPI.vl.imv.r<br>CPc.i.vm |  |  |  |  |  |
| 9 | CPI.vl.v.r<br>CPI.vl.vt (m/o) | CPI.vl.v.r<br>CPI.vl.vt (m/o) | CPI.vl.vt (m/o) | CPI.vl.vt (m/o) |  |  |  |
| 10 | CPr.l.vm<br>CPI.vm.v<br>CPc.v.vm<br>ACBsh.l<br>ACBsh.m | CPr.l.vm<br>CPI.vm.v<br>ACBsh.l | CPI.vm.v<br>CPc.v.vm<br>ACBsh.l<br>ACBsh.m |  |  |  |  |
| 11 | CPc.v.vm<br>CPext.cv | CPc.v.vm |  | CPc.v.vm | CPc.v.vm | CPc.v.vm<br>CPext.cv |  |
| 12 | CPr.imv<br>CPI.dm.cd<br>CPI.dm.im |  | CPr.imv<br>CPI.dm.cd<br>CPI.dm.im |  |  |  |  |
| 13 | CPr.l.ls<br>CPr.l.vm |  | CPr.l.ls<br>CPr.l.vm |  |  |  |  |
| 14 | CPI.vl.cvl<br>CPc.d.vm |  | CPI.vl.cvl<br>CPc.d.vm |  |  |  |  |
| 15 | CPI.dl.imd (ll)<br>CPI.vl.imv.r |  | CPI.dl.imd (ll)<br>CPI.vl.imv.r |  |  |  |  |
| 16 | CPI.vl.imv (ul) |  | CPI.vl.imv (ul) | CPI.vl.imv (ul) | CPI.vl.imv (ul) |  |  |
| 17 | CPI.vl.v (m/i)<br>CPI.vl.v.r |  | CPI.vl.v (m/i)<br>CPI.vl.v.r | CPI.vl.v (m/i)<br>CPI.vl.v.r | CPI.vl.v (m/i) |  |  |
| 18 | CPr.imd<br>CPr.l.ls<br>CPI.vm.v<br>CPI.vm.vm |  |  | CPr.imd<br>CPr.l.ls<br>CPI.vm.v<br>CPI.vm.vm |  |  |  |
| 19 | ACBsh.l |  |  | ACBsh.l |  |  |  |
| 20 | CPr.imv<br>CPI.vm.cvm<br>CPc.i.vm |  |  | CPr.imv<br>CPI.vm.cvm<br>CPc.i.vm |  |  |  |
| 21 | CPI.dm.cd |  |  | CPI.dm.cd |  |  |  |
| 22 | CPc.d.vm |  |  | CPc.d.vm |  |  |  |
| 23 | CPI.dl.d.r<br>CPI.dm.dl |  |  | CPI.dl.d.r<br>CPI.dm.dl |  |  |  |
| 24 | CPI.vl.cvl<br>CPc.d.dl |  |  | CPI.vl.cvl<br>CPc.d.dl |  |  |  |
| 25 | CPI.dl.d (tr)<br>CPI.dl.imd (ll)<br>CPI.vl.imv.r<br>CPc.i.d<br>CPc.i.vl |  |  | CPI.dl.d (tr)<br>CPI.dl.imd (ll)<br>CPI.vl.imv.r<br>CPc.i.d<br>CPc.i.vl |  |  |  |

Table 2. continued...
| GPe domains | Striatal inputs to<br>GPe domain | GPe levels |  |  |  |  |  |
| --- | --- | --- | --- | --- | --- | --- | --- |
|  |  | ARA 58 | ARA 60 | ARA 62 | ARA 64 | ARA 66 | ARA 68 |
| 26 | CPI.dm.dm<br>CPc.d.dl<br>CPc.d.dm<br>CPc.i.d |  |  |  | CPI.dm.dm<br>CPc.d.dl<br>CPc.d.dm<br>CPc.i.d |  |  |
| 27 | CPI.dl.d (tr)<br>CPI.dm.dl<br>CPI.vl.cvl<br>CPext.d |  |  |  | CPI.dl.d (tr)<br>CPI.dm.dl<br>CPI.vl.cvl<br>CPext.d |  |  |
| 28 | CPI.dl.imd (II) |  |  |  | CPI.dl.imd (II) |  |  |
| 29 | CPc.i.vl |  |  |  | CPc.i.vl |  |  |
| 30 | CPr.l.vm<br>CPI.vl.vt (m/o)<br>CPc.v.vl<br>CPext.rv |  |  |  | CPr.l.vm<br>CPI.vl.vt (m/o)<br>CPc.v.vl<br>CPext.rv | CPr.l.vm<br>CPc.v.vl<br>CPext.rv |  |
| 31 | CPc.d.dm |  |  |  |  | CPc.d.dm |  |
| 32 | CPc.d.dl<br>CPc.d.vm<br>CPc.i.d<br>CPc.i.vm<br>CPext.d |  |  |  |  | CPc.d.dl<br>CPc.d.vm<br>CPc.i.d<br>CPc.i.vm<br>CPext.d |  |
| 33 | CPc.v.vm<br>CPext.cv |  |  |  |  |  | CPc.v.vm<br>CPext.cv |
| 34 | CPc.d.dm<br>CPc.d.vm<br>CPc.i.vm |  |  |  |  |  | CPc.d.dm<br>CPc.d.vm<br>CPc.i.vm |
| 35 | CPext.d |  |  |  |  |  | CPext.d |
| 36 | CPext.rv |  |  |  |  |  | CPext.rv |

#### Convergent terminal fields in the striatonigral pathway define a nigral domain structure

At a finer scale of resolution, many of the individual striatal projection pathways show confluence and overlap in their nigral terminal fields, such that a domain structure of multiple convergent terminal zones is imposed upon the SNr by the striatonigral projectome. Computational analysis of the entire striatonigral dataset at each level of the SNr revealed 14 domains across the whole SNr (Figure 2G-I, 3A; Supplementary Figure 1, 3). Most of the CP domains generate axonal projections along the entire rostrocaudal extent of the SNr (Supplementary Figures 1B, 2, 3; Table 1) (Gerfen, 1985), a property we observed even at the single-cell level using fluorescence micro-optical sectioning tomography (fMOST) (Figure 3B), and which others have seen using biocytin injections or palmitoylated-GFP expression (Levesque & Parent, 2005; Fujiyama et al. 2011). As a result, many of the domains in the SNr formed by these inputs span multiple levels of the SNr, i.e., the SNr domains have a three-dimensional shape.

**Figure 3.**
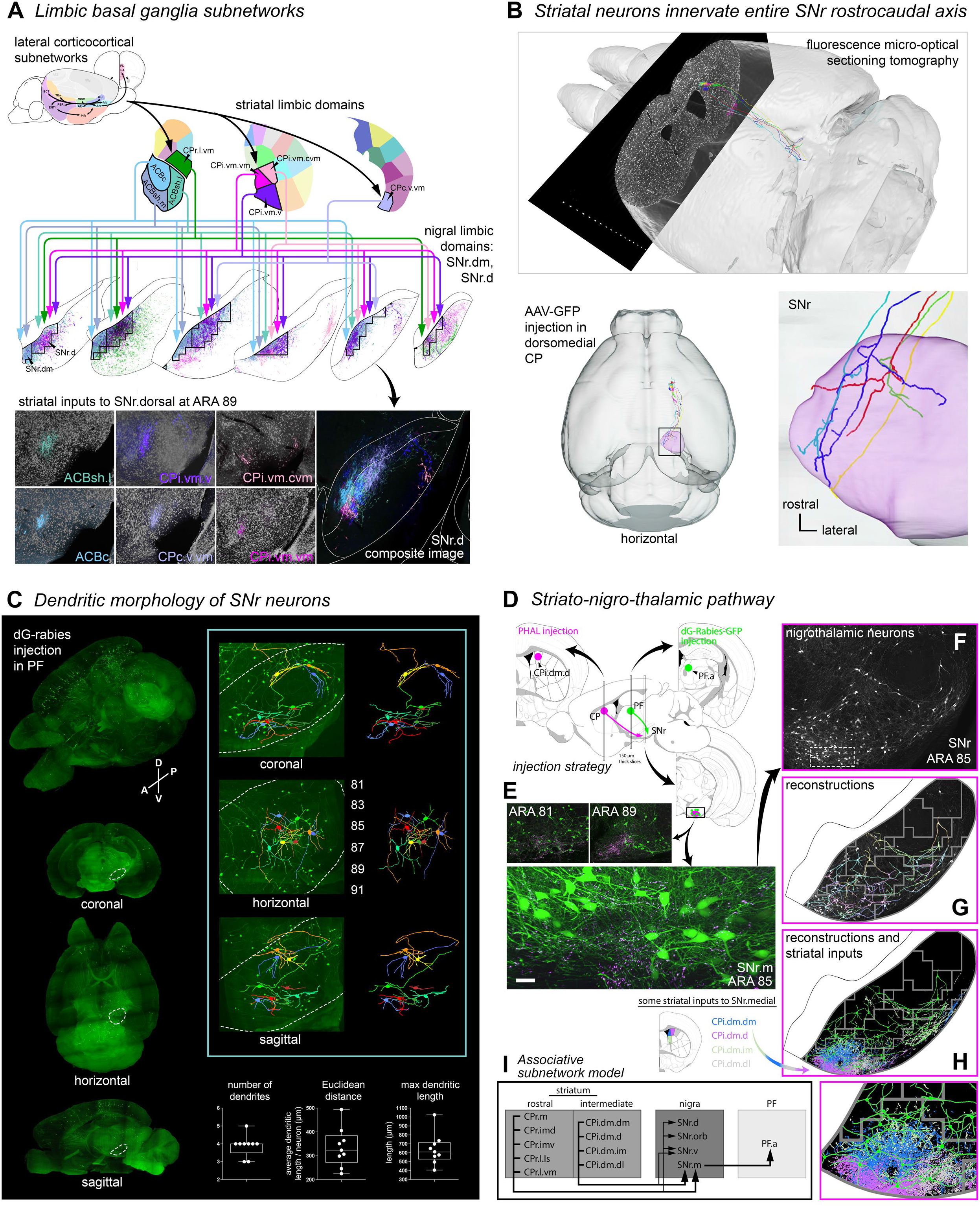
Convergent structure of the striatonigral pathway. A) Cortical limbic regions are networked together and they project to a common set of striatal domains; those domains project in complex convergent/divergent patterns to the SNr.dorsal (SNr.d) and SNr.dorsomedial (SNr.dm), forming the limbic cortico-striatonigral pathways. Raw axonal images, pseudocolored to match their striatal source domain, are shown individually and in composite (*bottom panels*). B) Individual striatal neurons were labeled with GFP and imaged with the fluorescence micro-optical sectioning tomography (fMOST) system, and many of them send axons that project through the rostrocaudal extent of the SNr. In total, 30 striatal neurons were reconstructed (Supplementary Figure 8), although only 5 are shown here for clarity. C) The dendritic morphology of SNr neurons was labeled with an injection of glycoprotein-deleted rabies in the parafascicular thalamus and digitally reconstructed. The whole-brain images show the labeling in different planes of view, and the SNr is outlined (dashed lines). The inset panel shows the SNr at high resolution, with the reconstructed neurons overlaid atop the images and alone on the right. The numbers beside the horizontal plane image indicate the approximate Atlas Level. The box plots at the bottom indicate some of the geometric features of the neurons (n=9), with boxes indicating 1^st^ and 3^rd^ quartiles and whiskers indicating min/max. D) The striato-nigro-thalamic pathway was labeled with an anterograde tracer injection in the CPi.dm.d and a retrograde tracer injection in the associative region of the PF (PF.a), and E) the labeling from both injections is seen in the SNr.medial (SNr.m) throughout its entire rostocaudal extent (scalebar = 30 µm). The nigrothalamic neurons at ARA 85 (*bottom image*) are shown in grayscale in (F). A sample of these neurons are reconstructed and shown in different colors in (G), while in (H) the same reconstructions are all colored green to differentiate them from the 4 axonal reconstructions of dorsomedial striatal inputs that terminate in the SNr.m (enlarged in the bottom image). This sample of neurons is reconstructed from a tissue slice 150 µm thick meaning that some dendrites were truncated, and thus the reconstructions best represent dendritic extent in the coronal plane. I) A wiring diagram of the associative subnetwork targeted in this experiment (not all connections are shown).

At the same time, the nigral terminations of a particular CP domain frequently shift in their mediolateral or dorsoventral position in the SNr at different rostrocaudal levels. Consequently, a single CP domain may contribute projections to multiple SNr domains, such as inputting a dorsal domain at one level and a medial domain at another. Most community detection algorithms impose a unitary structure on an information network such that each node belongs to one and only one community. This simplified network structure makes it easier to interpret, but may not be reflective of the complex nature of real world networks, wherein a single node may participate in multiple subnetwork structures, as is likely true in brain networks. We analyzed the striatonigral projection data level-by-level, and detected different network structures at each level of the SNr. We then manually joined similar communities across levels of the SNr, by grouping together communities that had similar input constituencies across levels of the SNr. For example, the striatal inputs to the medial region of the SNr are similar enough when comparing adjacent levels (mean Jaccard index comparing all adjacent levels=.61) that we determine them to form one continuous domain, the medial domain (Supplementary Figure 3, Supplementary Table 1). This approach resulted in 14 nigral domains, a number of which have 3-dimensional, cross-level structure (Figure 3A; Supplementary Figure 3). At the single neuron resolution (Figure 3B), we also demonstrated that axons arising from individual striatal projection neurons extend throughout the rostrocaudal length of the SNr and generate axonal branches.

There are several major domains that run the entire length of the SNr. The *medial* domain (SNr.m, see Figure 2I; Supplementary Figure 1B) is a highly integrative domain occupying the medial edge of the SNr, running its entire rostrocaudal length (Supplementary Figures 1B, 3). The SNr.m agglomerates inputs from most of the rostrocaudal axis of the CP: the CPr (CPr.m, CPr.imd, CPr.imv, CPr.l.ls, CPr.l.vm), most of the CPi.dm (CPi.dm.dm, CPi.dm.im, CPi.dm.d, and CPi.dm.dl) and even from the CPc (CPc.d.dm). Caudally at ARA 91 the SNr.m also incorporates inputs from the striatal trunk region (CPi.dl.d (tr)) as well as ACBsh.l. Although the roster of inputs to the SNr.m varies from level to level, the CPr.m, CPi.dm.im, and CPi.dm.d are inputs at every level (Figure 2E,G,H; Supplementary Figure 3; Supplementary Table 1).

The *dorsal* domain (SNr.d, see Figure 2I; Supplementary Figure 1B) receives inputs from many of the limbic domains of the CP: the CPi.vm.vm, CPi.vm.v, CPi.vm.cvm, CPc.v.vm, and CPr.l.vm, along with the lateral accumbens shell, ACBsh.l, and at caudal levels the core (ACBc) and medial accumbens shell (ACBsh.m) (Figure 2E,G,H; Supplementary Figure 3). The analysis grouped the CPi.dm.cd in the SNr.d at ARA 89, but we manually removed it because its terminations appeared too diffuse to properly characterize its location. The *dorsomedial* domain (SNr.dm) is a small region that receives a dense, concentrated input from the medial ACB, specifically the ACBc and ACBsh.m. It spans only the rostral half of the SNr, ARA 81-85, and at ARA 87 merges with the SNr.d (Figure 3A; Supplementary Figure 3). These contributory ACB and CP domains receive different combinations of inputs from the lateral cortico-cortical subnetworks (Hintiryan, Foster et al. 2016) whose component cortical regions are classically associated with the limbic system and emotion, specifically AId, AIp, AIv, PERI, PIR, VISC, ILA, PL, and TEa, as well as lBLAa, and all of that information converges in the SNr.d (Figure 3A).

The *oro-brachial* domain (SNr.ob) is the largest of the motor domains, and mainly integrates inputs from the mouth areas of the CP (CPi.vl.v (m/i), CPi.vl.vt (m/o), CPc.v.vl), the upper limb area (CPi.vl.imv (ul)), along with the CPi.vl.cvl and several limbic- and motor-related regions of the rostral and caudal CP (CPr.l.vm, CPc.i.d, CPc.i.vl, CPc.v.vl, CPc.v.vm) (Figure 2E,G-I; Supplementary Figure 3; Table 1). The terminal fields from CPi.vl.imv (ul) and CPi.vl.v (m/i) overlap extensively at every level of the SNr.orb except the rostral-most level, which receives only upper limb input.

The *lateral* domain (SNr.l) receives a mix of inputs from the domains of the CPc and the CPext. The *ventral* domain (SNr.v) gets the most consistent input from the trunk area CPi.dl.d (tr) and CPc.d.dl, and spans every level except ARA 91. The remaining domains span fewer levels (Supplementary Figure 3; Table 1). All together, the cortico-striato-nigral pathways form 14 output channels.

#### The nigral neurons within a domain project to the same thalamic target

Next, we investigated whether all neurons within the same newly defined SNr domain project to the same target as a validation of the accuracy of their delineations. This is demonstrated with an injection of anterograde tracer in the striatal domain CPi.dm.d and retrograde tracer dG-rabies-GFP (glycoprotein-deleted rabies, incapable of transsynaptic spread) in the associative sub-region of the parafascicular thalamic nucleus PF.a (see section *Thalamic connectivity* below). It shows that the striatal domain CPi.dm.d projects throughout the entire rostrocaudal extent of the SNr.m, and GFP-labeled PF.a-projecting nigral neurons are likewise observed throughout the rostrocaudal extent of the SNr.m (Figure 3D,E). Thus the matching structure of the inputs and the outputs of the SNr support the notion of longitudinal domains as a functional unit.

#### Dendritic arbors of nigral neurons conform to the size of their domain in dorsoventral and mediolateral dimensions

We observe the dendritic structure of nigral neurons with an injection of dG-rabies virus into the parafascicular thalamus, one of the principal efferent targets of the SNr. SHIELD-clearing (Park et al. 2019) and lightsheet imaging of the intact brain reveals the structure of a subset of nigral neurons in the coronal, horizontal, and sagittal planes (Figure 3C). Reimaging at high resolution permits reconstruction and visualization of those neurons in situ (Figure 3C, *inset*). In the horizontal plane (Figure 3C, *inset, middle panel*) their dendrites can be seen spreading partway into adjacent rostral and caudal levels, but they do not span the entire rostrocaudal length of the SNr. Quantification of the morphological features of these neurons support this (Figure 3C, *bottom*). The neurons have an average of 4 primary dendrites (mean: 3.89; range: 3-5). The dendrites of each neuron have a mean Euclidean length of 335±27.24 μm (mean±SE, from soma to distal tip, ignoring tortuosity), and the longest dendrite of each neuron averages 639±58.63 μm; this gives the neurons a ‘radius’ capable of extending at least a couple of atlas levels rostral and caudal to the soma. Many of these neurons’ primary dendrites are angled in asymmetric geometries, rather than in a maximally extended tetrahedral form. Collectively these data suggest an input integration capacity roughly on par with the dorsoventral and mediolateral dimensions of the nigral domains, i.e., these neurons appear capable of integrating most inputs to their domain at a given coronal level. However, their dendrites are not long enough to integrate inputs across the entire rostrocaudal length of the large domains, meaning that the small changes in striatal input structure across a domain at different rostrocaudal levels could result in neurons with slightly different integration profiles along the length of an SNr domain.

#### Neurons within a nigral domain are capable of integrating all inputs contributing to that domain at a given coronal level

In Figure 3H, manually traced graphic reconstructions of rabies-labeled SNr.m neurons projecting to the PF.a are shown overlaid atop a composite projection map of four of the striatal direct pathway inputs to that nigral domain. Those striatal terminal fields topographically innervate separate but overlapping subregions within the SNr.m, while the dendrites of most of the postsynaptic SNr.m neurons extend sufficiently that they can receive contacts from all of the striatal terminal fields. Furthermore, even though recipient neurons within the domain are physically capable of integrating all inputs, they likely have heavier inputs from one input source based on the differing densities of axonal terminals within their dendritic arbors. This could have important functional implications for the nigral neurons, because the structure of this pathway appears to permit both a convergence of many striatal inputs onto individual nigral neurons, as well as a bias toward one or a few particular inputs over the others. Also, it can be seen that dendrites of neurons near the margins of a domain often (though not always) extend into adjacent domains, indicating that they could participate in a functional crossover at domain borders; this kind of overlap at the margins of functional domains is seen elsewhere in the basal ganglia (Hintiryan, Foster et al. 2016; Haber, 2016), and may be an important property to allow integration across information streams.

### Organizational principles of the striatopallidal pathway

Three additional striatal injection sites were included in the GPe analysis (see Methods). These three were from a level of the CP in between the CPr and CPi. They were included because they labeled zones in the GPe that were not targeted by any other injection site’s terminals. This is remarkable because it reveals that the GPe has a much more faithful topography of the CP than does the SNr (elaborated below).

#### The striatopallidal projectome is typified by a high degree of parallelism

The striatopallidal indirect pathway is characterized by a higher degree of parallelism than the striatonigral pathway, with far less convergence of inputs arising from different regions of the striatum. In general, most domains of the CP project to their own unique area in the GPe with little overlap between adjacent terminal fields. The largely parallelized striatopallidal topography is evident at the division, community, and domain level in the reconstruction maps. At the division level, the CPr projects to rostral GPe, the CPi to intermediate levels of the GPe, the CPc to caudal levels of the GPe, and the CPext to the most caudal levels of GPe, a topography similar to the primate striatopallidal pathway (Hazrati & Parent, 1992) (Figure 4A, Supplementary Figure 2B). The community-level projections illustrate that the pallidal terminal fields maintain the dorsoventral and mediolateral topographic relationships of their striatal origins (Figure 4B). In the domain-level reconstruction maps, each striatal terminal field is colored according to its striatal source domain (Figure 4C). Many more individual colors can be seen compared to the nigral domain-level reconstruction maps (Figure 2E), an indication of the parallelization of the striatopallidal pathway compared to the striatonigral pathway, which shows more overlap and blending of the colored terminal fields. The pallidal somatotopic body map runs along the lateral edge of the GPe at ARA 60 and 62 (Figure 4D) and follows the same tail up, head down orientation of the striatal body map (GPe domains 6, 9, 15-17, and 25, see next section).

**Figure 4.**
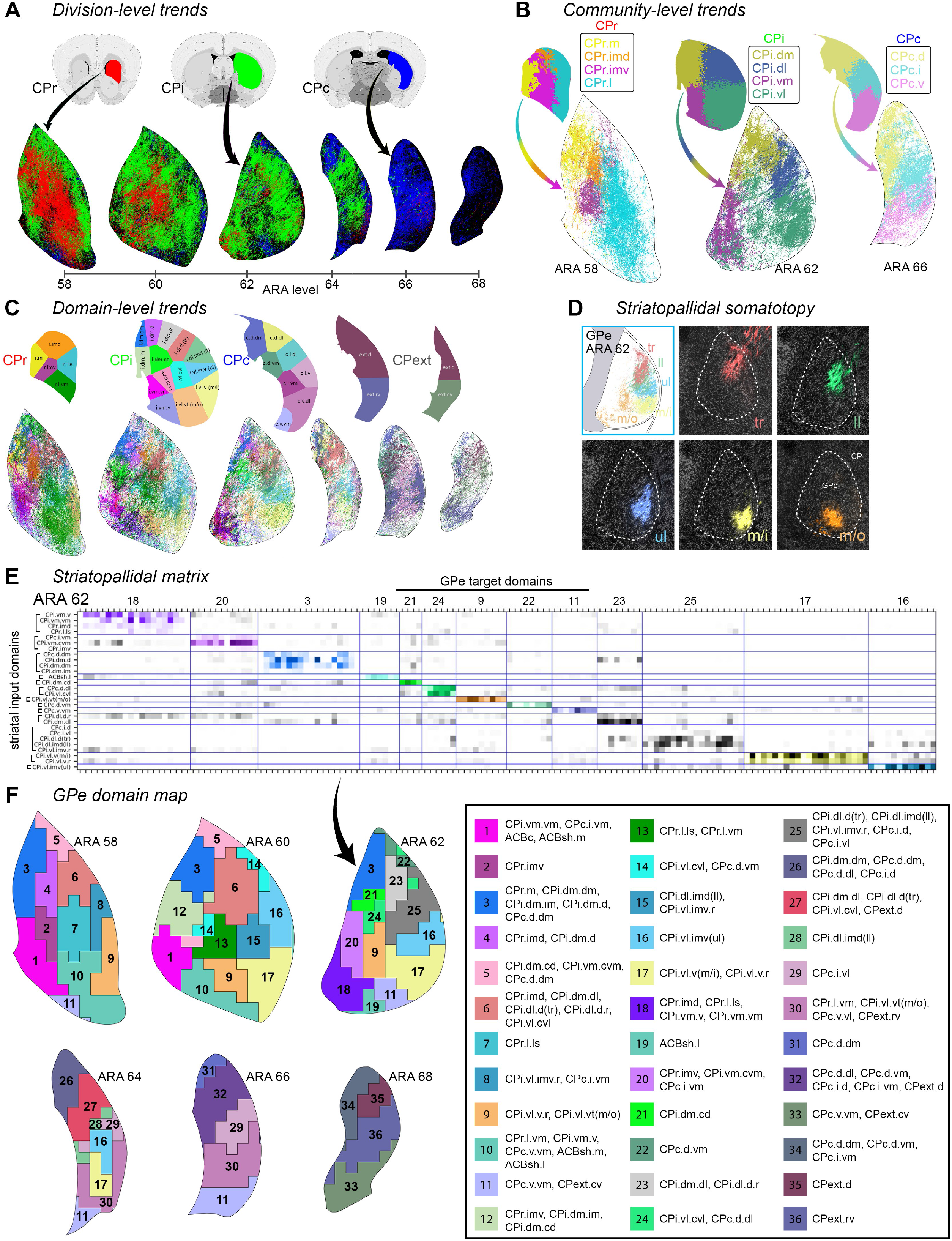
Projection trends of the striatopallidal pathway at different scales and novel GPe domain map. A) Axonal reconstructions originating from all striatal domains of each CP division are colored red, green, or blue, accordingly. The rostrocaudal axis of the CP terminates in a rostrocaudal pattern in the GPe, respectively. B) Community-level view of striatopallidal terminations depicts axons of all CP domains colored to match their parent community (e.g., axons of injections in CPi.dl.d and CPi.dl.imd are colored blue for community CPi.dl), revealing that the dorsoventral and mediolateral topography of the CP is generally maintained in the GPe. C) Domain-level striatopallidal terminations are colored according to their CP source domain. D) The somatotopic map in the GPe is revealed by striatopallidal terminations from body region-specific striatal source domains. E) The striatopallidal matrix for GPe at ARA 62. Note how each domain has fewer inputs than the nigral domains (Figure 2G, Supplementary Figures 1, 4), and even the pallidal domains with 4 or 5 inputs tend to be densely innervated by only 1 or 2 of those inputs. The arrow points to its map in (F). F) The novel GPe domain map: domains are input-defined by the striatopallidal projectome (see figure legend).

#### Striatopallidal terminal patterns define a pallidal domain structure

Computational analysis of the striatopallidal axonal data reveals a high dimension, variegated domain structure in GPe (Figure 4E,F, Supplementary Figure 4, Supplementary Table 2). In all, the striatopallidal projectome partitions the GPe into 36 domains (Figure 4F). The majority of domains (64%) have only one or two inputs, and most domains (69%) only span one coronal level of the GPe. This reflects the specificity and parallelization of the striatopallidal pathway compared to the striatonigral pathway, and indicates a higher degree of faithful transposition of striatal information into the GPe, with less convergence and funneling of that information. Further, even though about one-third of GPe domains receive 3 or more inputs, most of these domains tend to have 1-2 dominant inputs, with the remaining inputs contributing substantially fewer axons.

One of the notable exceptions to these parallelization trends is the domain GPe.3 (Figure 4F). This domain is one of the few GPe domains that receives multiple inputs and extends through several of the rostrocaudal levels of the GPe. Interestingly, it is similar to the SNr.m in that all of the inputs that group together in GPe.3 (i.e., CPi.dm.dm, CPi.dm.im, CPi.dm.d, CPr.m, and CPc.d.dm) also group together in SNr.m, and in fact are among the densest and most consistent inputs to SNr.m. These striatal regions comprise the dorsomedial corner of the CP adjacent to the lateral ventricular wall all along its rostrocaudal length (Figure 4C). They collectively receive a highly similar set of cortical inputs as well (Hintiryan, Foster et al. 2016), cortical areas that themselves form the two medial sensory associative cortico-cortical sub-networks (Zingg, Hintiryan et al. 2014). Thus the two sensory associative cortical sub-networks project to a small set of striatal domains, which in turn send convergent projections to domains in the GPe and SNr.

#### The direct and indirect pathways do not have topographic differences in GPe

The direct (striatonigral) and indirect (striatopallidal) pathways differ anatomically in that the indirect pathway projects only to GPe while the direct pathway is assumed to project only to the SNr/GPi. However, single neuron tracing studies have shown that nearly all direct pathway neurons send a collateral axon to the GPe as they course to the SNr/GPi (Kawaguchi et al. 1990; Wu et al. 2000; Levesque & Parent, 2005; Fujiyama et al. 2011), an offshoot termed a bridging collateral (Cazorla et al. 2014). Although the direct and indirect pathway populations are intermingled in the CP (Flaherty & Graybiel, 1993) it is unknown if their topography differs. A rich knowledge of the topography of these two pathways is critical for assembling detailed network schematics and understanding their functionality. Thus we injected adenovirus expressing Cre-dependent GFP or RFP into the CPi.dm and CPi.dl of D1-Cre (for direct pathway) and A2A-Cre (for indirect pathway) strains of mice. The results reveal that both direct and indirect pathway neurons originating in the same region of the CP target the same terminal region of the GPe (Figure 5A). Although not quantified, the density of the indirect pathway terminations in GPe appears denser than the direct pathway. Tracing data using the fMOST technique (Figure 5B) also shows at a single-neuron level that indirect pathway neurons (red, orange) have a denser pallidal terminal field than the bridging collateral terminal fields of the direct pathway (blue, green, aqua), though they do have similar topography. Additionally, the results confirm that the indirect pathway projects to GPe only while the direct pathway projects to both GPe and SNr/GPi (Cre-dependent data not shown).

**Figure 5.**
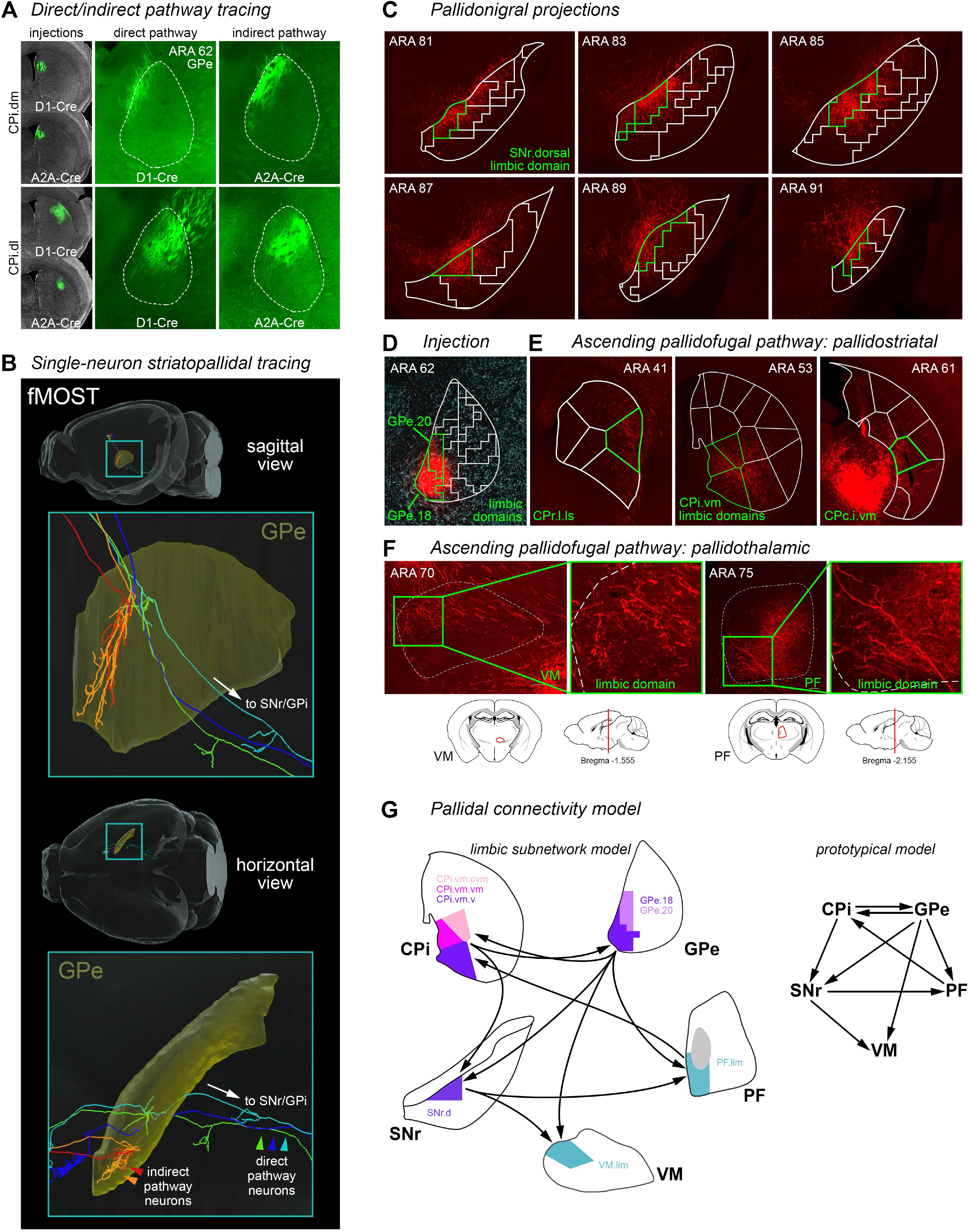
Striatopallidal and pallidofugal connectivity. A) Direct and indirect pathway striatopallidal topography was compared by Cre-dependent tracing of D1 dopamine receptor-expressing (direct) and A2A adenosine receptor-expressing (indirect) medium spiny neurons. Injections of AAV-DIO-EGFP were made into the same region of dorsomedial (CPi.dm) or dorsolateral (CPi.dl) striatum in both mouse lines (n=2 each). Direct and indirect pathway terminals from the same striatal region had the same topography in the GPe. The indirect pathway terminal fields were consistently denser than those of direct pathway, even though in one case (CPi.dl), the direct pathway injection was clearly larger. These properties are also borne out at the single-cell resolution as seen in the fMOST data in (B), where 2 indirect pathway neurons (red and orange) terminate more extensively than 3 direct pathway neurons (blue, green, and aqua), while they all terminate in the same general region (also see Supplementary Figure 8). C) The GPe in turn projects densely to the SNr with a homotypic topography. In this case, an injection in two of the GPe limbic domains (D, GPe.18 and 20, highlighted green) labels axonal terminals in the nigral limbic domain SNr.dorsal (highlighted green in [C]). The SNr.dorsal, GPe.18, and GPe.20 are all innervated by the same striatal domains. The same pallidal injection also labels axons that project: E) back into the striatum, reciprocally innervating the input striatal domains (highlighted green); and F) to the limbic domains of the ventromedial (VM) and parafascicular (PF) thalamic nuclei. G) All of this connectivity is summarized in the limbic subnetwork and prototypical wiring diagrams.

#### The pallidonigral pathway projects along the entire rostrocaudal length of the nigra, and it converges with the striatonigral pathway in a homotypic fashion

The GPe completes the indirect pathway by projecting to the SNr (and subthalamic nucleus, not discussed here) (Smith & Bolam, 1989). Even though striatal projections to the GPe innervate only a restricted rostrocaudal range of the pallidum (Figure 4A, 5B), projections from GPe to nigra innervate its entire rostrocaudal length in a topographic fashion (Figure 5C), similar to the striatonigral pathway (Figure 2A,B).

Moreover, the pallidonigral projections from a GPe domain converge with striatonigral axons arising from the same CP domain that serves as input source to both nuclei. Direct and indirect pathway neurons are intermingled in the striatum, such that one domain of the CP projects to specific domains of the GPe and SNr (Figures 2-4). We find that the GPe domain targeted by the indirect pathway projects to the nigral domain targeted by the direct pathway, forming a specific indirect pathway subnetwork. For example, an anterograde tracer injection in the limbic GPe domains 18 and 20 (Figure 5D) labels projections to the nigral limbic domain SNr.d (Figure 5C, SNr.d is highlighted green). Both the GPe.18 and 20 and the SNr.d receive inputs from striatal domains CPi.vm.v, CPi.vm.vm, and CPi.vm.cvm. Thus the direct and indirect pathways arising from a common striatal source converge in the SNr. In accord with this, a previous electron microscopy experiment found synaptic terminals from both pathways on individual SNr neurons (Smith & Bolam, 1991). It has also been found that during behavioral performance both the direct and indirect pathways must be activated for appropriate behavioral responding (Cui et al. 2013; Tecuapetla et al. 2016). Our data describes an anatomical substrate for the coordination of those two pathways during behavior.

#### The pallidostriatal pathway projects topographically back to the CP domains that innervate it

The pallidofugal pathway contains an ascending portion that projects back to the striatum. Many prototypic pallidal neurons that project downstream to the SNr, STN, and/or GPi also send a collateral up to the CP, and a unique population of neurons called arkypallidal neurons richly project back to the CP and to no other target (Sato, et al. 2000; Mallet et al. 2012; Mastro, et al. 2014; Abdi, et al. 2015; Simmons, et al., 2020). However, no specific topography has been described for this pathway. We find that the pallidostriatal projection from a GPe domain largely projects back to the same CP domains that provide input to that GPe domain. From the same injection in pallidal domains GPe.18 and 20 used to demonstrate the pallidonigral pathway (Figure 5D), pallidal axons can be seen terminating in CPr.l.ls, CPi.vm.v, CPi.vm.vm, CPi.vm.cvm, and CPc.i.vm (Figure 5E, relevant domains are highlighted green), all of which are striatal domains that generate inputs to GPe.18 and 20 (Figure 4F).

#### The pallidothalamic pathway projects directly to PF and VM with topographic specificity

The GPe projects directly to the thalamic nodes of the cortico-basal ganglia loop, bypassing the traditional basal ganglia output nuclei of the SNr/GPi. From the same injection demonstrating the pallidonigral and pallidostriatal projections (Figure 5D), labeled pallidal axons can be clearly seen terminating in the dorsomedial zone of the VM and the ventromedial zone of the PF (Figure 5F), both of which are the limbic sub-regions of those nuclei (see next section, *Thalamic connectivity*). Those thalamic regions both receive input from the nigral limbic domain SNr.d (ibid.). Previous work demonstrated projections from GPe to PF (Mastro et al. 2014; Saunders et al. 2015), but we present the first demonstration of GPe projections to VM, as well as the topographic specificity of both pallidothalamic pathways. Thus the GPe is itself a striatal output region and it is specifically interconnected with all other homotypic subregions in the basal ganglia-thalamic subnetwork (Figure 5G).

### Thalamic connectivity

The output nuclei of the basal ganglia, the SNr and GPi, send projections to a number of thalamic nuclei, such as parafascicular (PF), ventromedial (VM), mediodorsal, and the ventral anterior-lateral complex, which in turn project back to cortex to complete the cortico-basal ganglia-thalamocortical loop. Some of the densest nigrothalamic projections are to the PF and VM, and here we reveal novel organizational principles of these nuclei as output channels for basal ganglia outflow.

#### Output channels of the PF

In the PF we examined the distribution patterns of corticothalamic and nigrothalamic axonal terminations, as well as patterns of thalamocortical and thalamostriatal cell groups. Remarkably, cortical areas, striatal domains, and nigral domains that are interconnected all have convergent input/output connectivity patterns with discrete zones of the PF, forming a unique subnetwork (Figure 6C), similar to patterns described for cortico-thalamo-striatal networks (Mandelbaum, et al., 2019). Based on these patterns, we identify 6 subdivisions of the PF which serve as parallel output channels for integrating basal ganglia efferent signals with cortical inputs, and conveying that computation to striatum and cortex (Figure 6A). Radiating out around the fasciculus retroflexus, these 6 regions correspond to associative (PF.a), trunk/lower limb (PF.tr/ll), upper limb (PF.ul), mouth (PF.m), limbic (PF.lim), and ventral striatal (PF.vs) channels. As an example, the associative subnetwork (Figure 6D,E, *top*) contains the dorsal prelimbic and frontal eye field cortical regions, which project to the CPi.dm.im and CPi.dm.d, respectively, and also to the PF.a; those striatal domains both receive input from the PF.a, and they both project to the SNr.m, which in turn projects to the PF.a; the PF.a projects back up to those cortical regions, closing the circuit. The 6 PF regions exhibit a gross topographical relationship with the major community-level regions of the striatum (Figure 6B). Collectively, the 6 output channels in the PF are organized with specific cortical, thalamic, and nigral subregions into 6 parallel cortico-basal ganglia subnetworks (Figure 6E).

**Figure 6.**
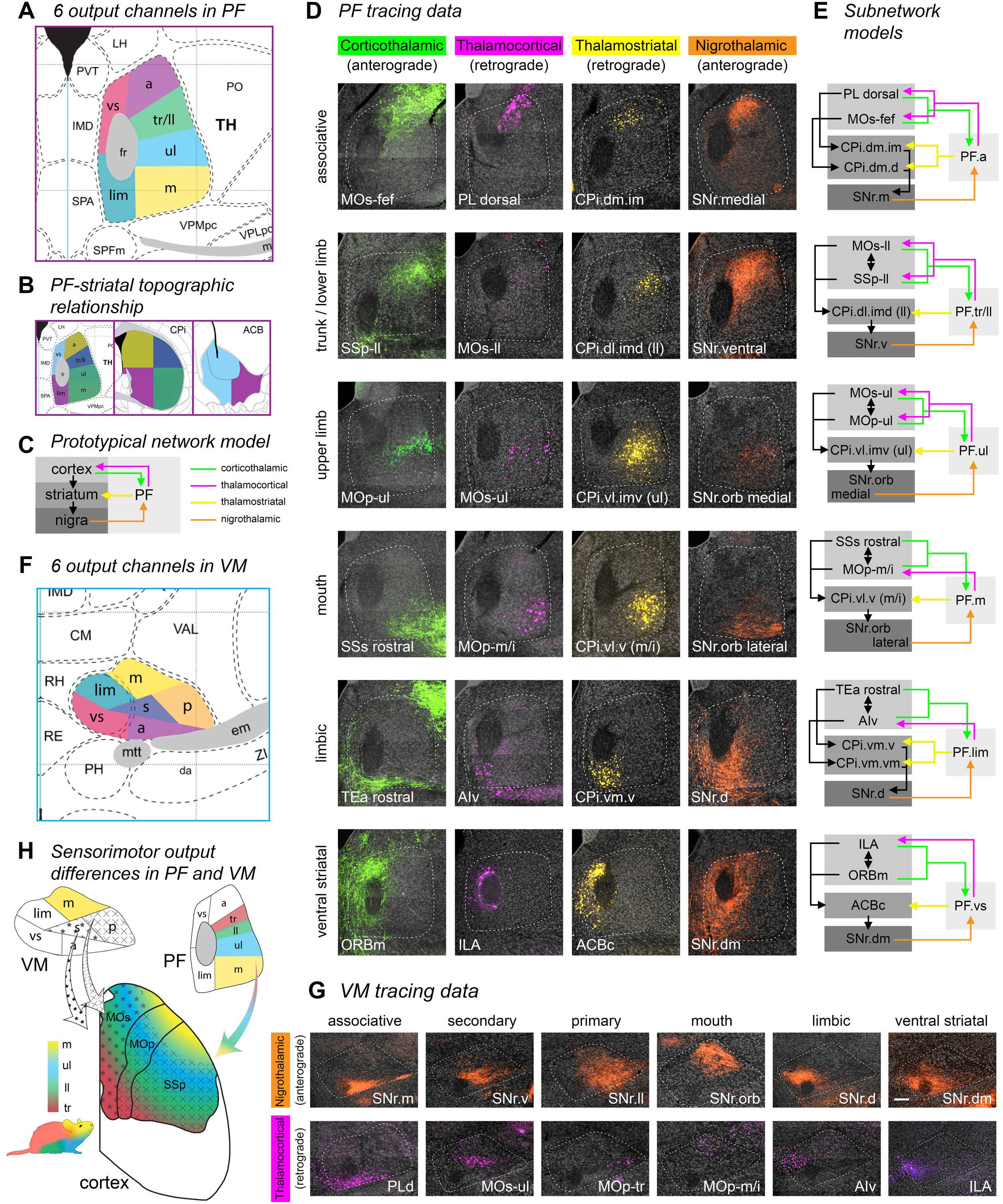
Thalamic connectivity. A) The parafascicular thalamus (PF) has 6 output channels, territories that are interconnected with domains in the basal ganglia and cortical regions that are themselves interconnected; the colored maps in (B) depict the general PF-striatal connectivity relationships. C) The prototypical network model illustrates the basic connectivity scheme of the cortico-basal ganglia-thalamocortical circuit. D) The connectivity of each of the 6 PF output channels with every other major node of the cortico-basal ganglia network (except GPe) is shown. The labeling for each pathway is pseudocolored to match the color scheme in (C). The nigrothalamic pathway data in the ventral striatal subnetwork (i.e., the image labeled ‘SNr.dm’) was generated using anterograde transsynaptic tracing, and thus represents synapse-specific pathway tracing from ACBc to SNr.dm to PF.vs (see main text for details). In (E) the specific subnetwork models are shown to the right of each corresponding image series. F) The ventromedial thalamic nucleus (VM) also has 6 output channels, but in VM the motor output pathways to cortex are not organized somatotopically by body subregion, as they are in PF, but rather the VM.primary (VM.p) projects to all body regions of primary motor and primary somatosensory cortex, and the VM.secondary (VM.s) projects to all body regions of secondary motor cortex. The limbic (lim), ventral striatal (vs), associative, and mouth (m) output channels are similar to the corresponding channels in PF. G) Nigrothalamic and thalamocortical tracing data supports the 6 channel model of VM. Note that the same transsynaptic tracing experiment described for (D) also resulted in the ventral striatal data in (G) (i.e., the image labeled ‘SNr.dm’). H) A summary diagram illustrating the thalamocortical projection differences and similarities between VM (shapes) and PF (colors). The tr and ll zones are separated in this PF model because tr and ll exhibit a partially overlapping, partially separable thalamocortical connectivity pattern (data not shown), and emphasizing them separately underscores the somatotopic features of this pathway. Scalebar = 50 µm.

To show the actual synaptic specificity of one of these pathways, the ACB-SNr.dm-PF.vs pathway of the ventral striatal subnetwork (Figure 6E, *bottom*), we used the anterograde transsynaptic tracing technique (Zingg et al. 2017). This method leverages the fact that when AAV1 is injected at sufficiently high concentration into a neuronal population, viral particles will travel down the axons and be released from the terminals where they can infect postsynaptic neurons (Zingg et al. 2020). Transsynaptic infection is an inefficient process and the postsynaptic viral load is comparatively low; but when paired with a highly sensitive technique like the Cre-Lox system, where very little Cre is required to unlock a Floxed gene like Flex.RFP, the transsynaptic infectivity rate is sufficient for tracing pathways. Not only does this combinatorial approach demonstrate synaptic specificity in the pathway traced, it can also be utilized to precisely label small targets like the SNr.dm. We injected AAV1.hSyn.Cre.WPRE.hGH into the ACBc and the Cre-dependent tracer AAV1.CAG.Flex.tdTomato.WPRE into the medial SNr at ARA 81-85. Labeled neurons are seen specifically in the SNr.dm at those levels (Supplementary Figure 7), and the axonal labeling from this nigral domain terminates precisely in the predicted regions of the PF and VM where the other nodes of the ventral striatal subnetwork connect, the PF.vs (Figure 6D, *bottom*) and VM.vs (Figure 6G, *right*). While neurons in the SNc and VTA were also labeled in this experiment, they do not contribute to the labeling seen in either the PF or VM, because staining for tyrosine hydroxylase reveals little to no catecholaminergic neurites in either thalamic nucleus (data not shown).

#### The VM thalamic nucleus has a different input-output organization than PF

The SNr also projects to the VM and adjacent submedial nucleus of the thalamus (SMT), and its outputs to that region are even denser than to the PF. Like the PF, the VM also appears to host at least 6 output channels (Figure 6F,G), but with a different organizational principle than the PF. Unlike the PF, with its separated homunculus of output channels for the body sub-regions, the VM has one output channel projecting to secondary motor cortex (VM.s) and one projecting to primary motor and primary somatosensory cortex (VM.p). The VM.s output channel projects to all somatic subregions (i.e., ul, ll, and tr) within the secondary motor cortex (MOs), while the VM.p projects to all somatic subregions (ibid.) within the primary motor (MOp) and primary somatosensory (SSp) cortex (Figure 6G,H). A separate output channel for the mouth pathways, VM.m, projects specifically to the mouth and head regions of MOp, SSp, and MOs, an arrangement similar to the PF mouth output channel. The sensory associative (VM.a), limbic (VM.lim), and ventral striatal (VM.vs) channels are similar in structure to the homologous regions of the PF. The boundaries of each of these output channels are established by the position of the thalamocortical neuron groups (Figure 6G). Each of these VM domains receives a specific nigral input from at least one of the SNr domains, and that nigrothalamic input conforms to the VM domain boundaries established by the thalamocortical output channels (Figure 6G).

### The direct corticonigral projection

The most recent major modification to the classical model of basal ganglia network structure was the incorporation of the hyperdirect pathway (Nambu et al. 1996), a glutamatergic cortical input to the subthalamic nucleus (Nambu et al. 2000), which in turn sends its own glutamatergic input to the SNr/GPi (Robledo & Feger, 1990). This excitatory disynaptic pathway bypasses the direct and indirect pathways of the striatum and GPe, and was thought to be the shortest route for cortical information to reach the SNr (Nambu et al. 1996, 2002), causing an early excitatory response in SNr neurons that precedes the inhibitory response mediated by the striatal direct pathway (Maurice et al. 1999). In fact we find that a limited number of cortical areas send axon collaterals directly to the SNr where they make synaptic terminations.

#### Cortical areas of the oro-brachial subnetwork project directly to SNr

Cortical injections of anterograde tracers reveal fine axonal fibers within the SNr (Figure 7A). These corticonigral axons have the highly branched, irregular appearance of an axonal terminal field, compared to the more uniform, unbranched, fasciculated arrangement of fibers of passage. Only a subset of cortical areas participate in this pathway: the inner mouth, outer mouth, and upper limb cortical subnetworks that feed into the oro-brachial output channel preferentially target the caudal levels of the SNr.orb. Most other cortical regions lack direct nigral projections (data not shown). Additionally, not every cortical area participating in the oro-brachial network sends direct projections: we see projections from MOp-m/i, SSp-m/i, SSs rostral, MOp-m/o, MOs-m/o, SSs caudal, and MOs-ul; but not from MOs-m/i, SSp-m/o, MOp-ul, SSp-ul, or SSp-bfd. Thus each cortical subnetwork has a unique pattern of corticonigral projections.

**Figure 7.**
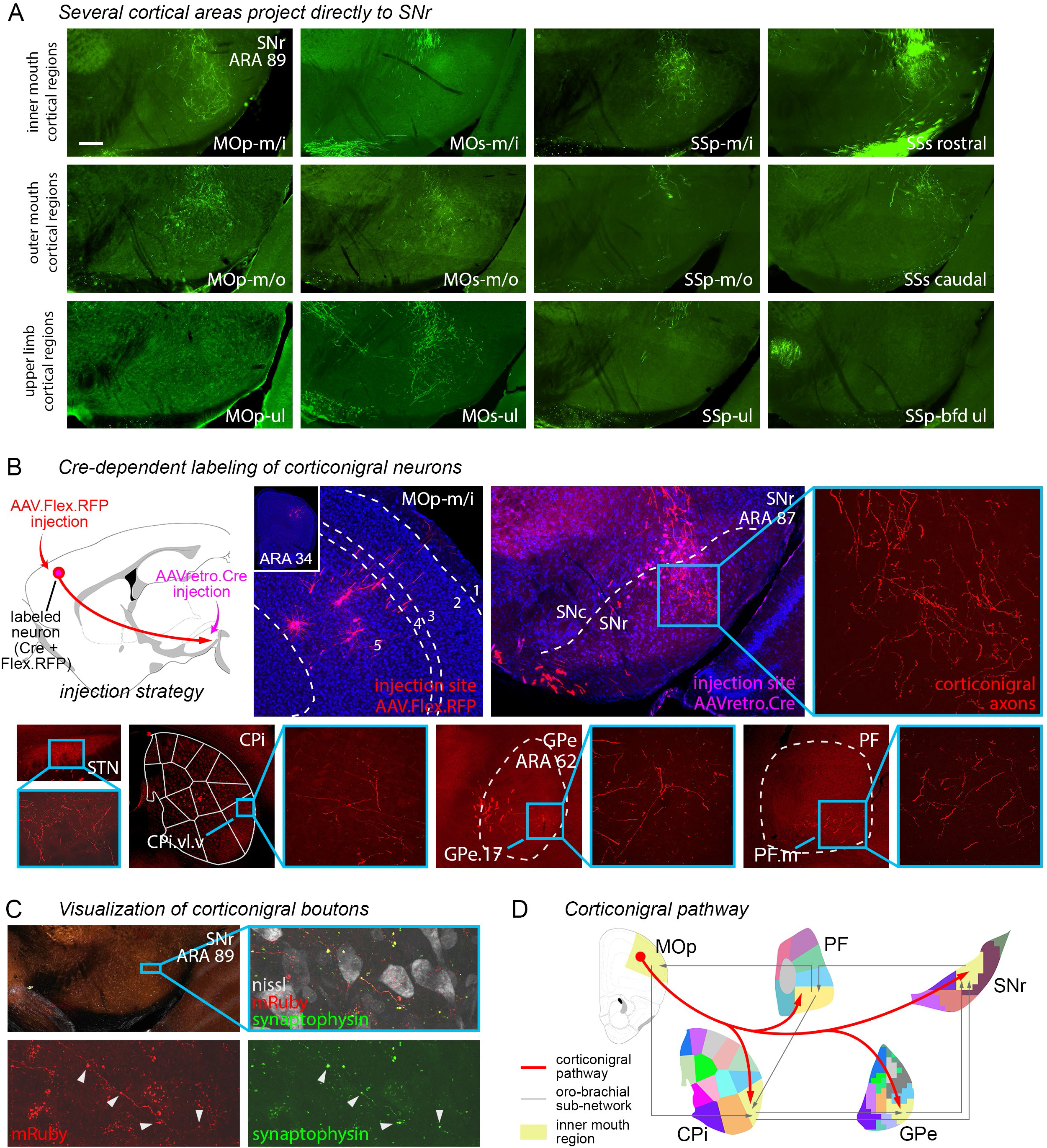
The cortico-nigral pathway. A) Some but not all cortical regions of the inner mouth (m/i), outer mouth (m/o), and upper limb (ul) somatomotor subnetworks send direct axonal projections to the SNr.oro-brachial domain, as demonstrated by PHAL injection. B) The pathway from m/i primary motor cortex (MOp-m/i) was validated using Cre-dependent labeling, by infecting cortical axons in the nigra with AAVretro.Cre, which unlocked fluorophore expression in MOp-m/i neurons injected with AAV.Flex.RFP. The labeled MOp-m/i axons are seen in the SNr.orb, as well as the mouth domains of the CP, GPe, and PF, as well as a restricted region of the STN, indicating that at least some of these cortical neurons comprise the hyperdirect pathway. C) To visualize bona fide synaptic terminations in the SNr, the MOp-m/i was injected with AAV.Syp-GFP.mRuby, which labeled axons with red fluorescence and synaptic boutons with green fluorescence (arrowheads). D) The corticonigral pathway appears to be a unique feature of the oro-brachial subnetwork, contacting all mouth regions in it and suggesting that cortex can simultaneously activate all other nodes.

#### Cre-dependent tracing confirms corticonigral pathway

It remains possible that these corticonigral collaterals are nonetheless non-terminating fibers of passage, so we implemented an intersectional approach to label them. First, we injected the SNr.orb with AAVretro.Cre, which retrogradely infects cortical neurons and induces Cre expression in them, while simultaneously injecting a Cre-dependent anterograde tracer (AAV1.FLEX.tdtomato) into the MOp-m/i cortex (Figure 7B). Again, we see labeled cortical axons impinge deeply into the SNr, and again we see a topographic specificity: the MOp-m/i projects directly to and terminates within the SNr.orb (Figure 7B). Axon collaterals from this injection also terminate within the STN, indicating that at least some of these fibers constitute part of the hyperdirect pathway (Figure 7B). Importantly, these corticofugal axons also send collaterals to the orofacial regions of every other important node of the orobrachial sub-network, specifically the CPi.vl.v (m/i), the GPe.17, and the PF.m (Figure 7B,D), as well as the VM.m (data not shown).

#### Corticonigral axons have terminal boutons in the SNr

To ascertain whether these corticonigral axons are genuinely making synaptic terminations in the SNr, we injected an AAV inducing expression of GFP-tagged synaptophysin and cytoplasmic RFP (AAV1.syp-GFP.mRuby). Since synaptophysin is an abundant integral membrane component of synaptic vesicles (Wiedenmann & Franke, 1985; Navone et al. 1986), its presence in the corticonigral collaterals would be strongly indicative of synaptic connectivity. Accordingly, injection of this virus into MOp-m/i labels red corticonigral axons in the SNr.orb bearing green terminal boutons (Figure 7C). This definitive demonstration and characterization of the corticonigral pathway suggests that the cortex can simultaneously activate virtually all components of the oro-brachial cortico-basal ganglia-thalamic subnetwork (Figure 7D).

### The cortico-basal ganglia-thalamo-cortical “loop”

The cortico-basal ganglia-thalamo-cortical loop has long been proposed as the core architectural form of neural networks involving the striatum, pallidum, and nigra (Alexander et al. 1986; Alexander et al. 1990; Parent & Hazrati, 1995; Swanson, 2000). However, calling the network a “loop” begs the question: Is it in fact a closed-circuit loop? The notion of the loop is supported by piecemeal electrophysiological and anatomical experiments that have demonstrated segments of these networks (Deniau et al. 1976; Deniau & Chevalier, 1985; Nambu et al. 1988, 1991; Hedreen & DeLong, 1991; Hoover & Strick, 1993; Rouiller et al. 1994; Kitano et al. 1998; Middleton & Strick, 2002; Tanaka et al. 2018). Yet entire intact circuitous loops have not been demonstrated within a single animal using any methodology. Nor has it been shown that the thalamocortical output actually feeds back onto the corticostriatal input neurons of that loop in a closed-circuit manner.

#### Anatomical demonstration of the complete oro-brachial subnetwork loop

Using the double co-injection technique for mapping interconnected network structures (Thompson & Swanson, 2010; Zingg, Hintiryan et al. 2014), we injected two anterograde/retrograde tracer pairs into two non-adjacent nodes of the oro-brachial cortico-basal ganglia subnetwork (i.e., either CP and PF, or CTX and SNr). In a serially connected 4-node loop, the labeling from the two injection pairs will converge and appose in the non-injected nodes (Figure 8A, *injection strategy*). Injections into the striatal and thalamic orofacial regions (CPi.vl.v (m/i) and PF.m, respectively) reveal overlapping labeling in the nigral domain SNr.orb and a cortical column in MOp-m/i (Figure 8A, *cortical labeling*). The same strategy also demonstrates the associative subnetwork loop (Supplementary Figure 6).

**Figure 8.**
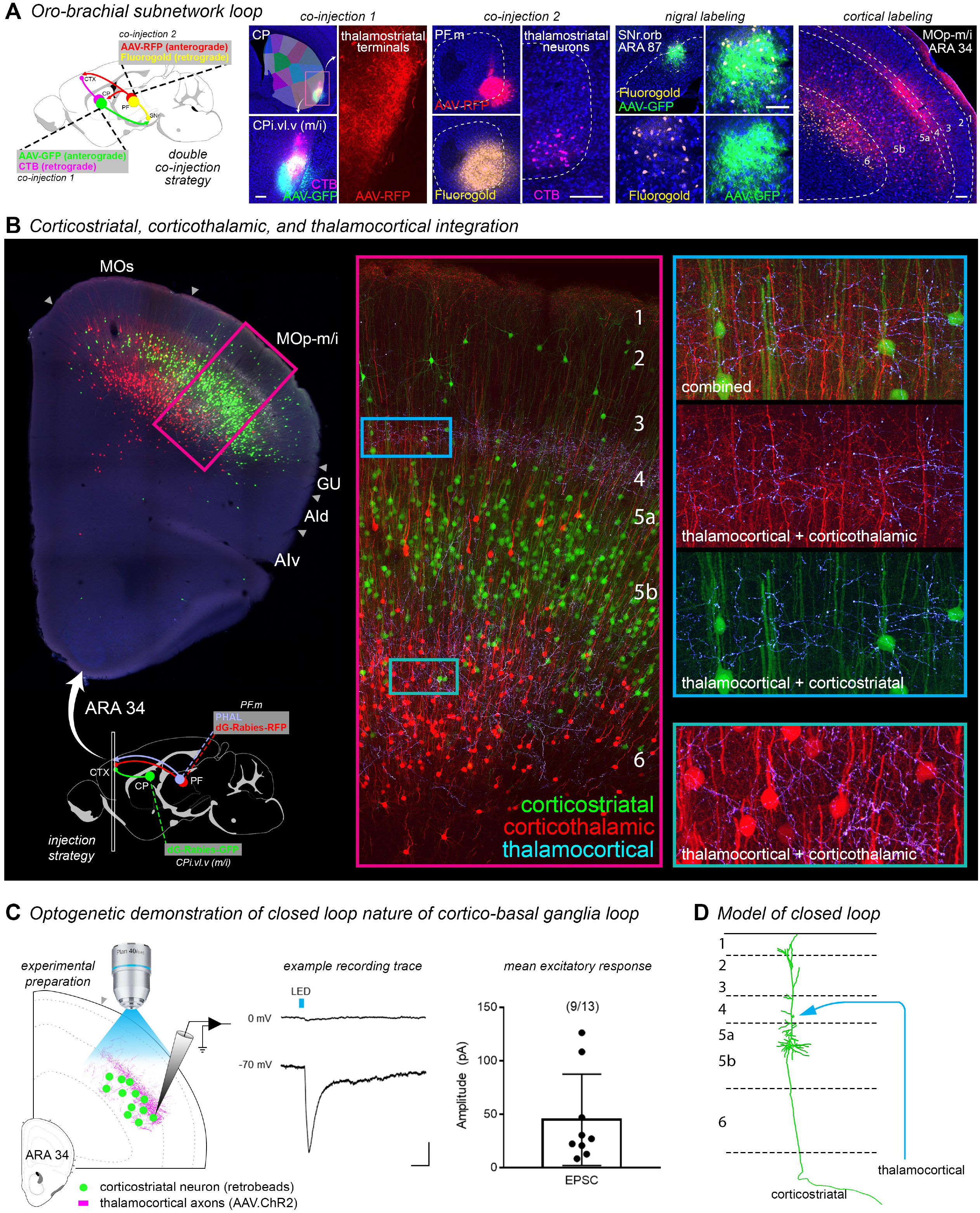
The cortico-basal ganglia-thalamo-cortical loop. A) The double co-injection technique was used to demonstrate complete cortico-striato-nigro-thalamocortical subnetwork loops, by injecting one co-injection (anterograde+retrograde tracer mix) into the striatal node and another co-injection into the thalamic node of a subnetwork. The transported tracers co-label precise subregions of the non-injected nodes (i.e., cortex and nigra). The photomicrographs demonstrate the oro-brachial subnetwork loop. B) To investigate how the thalamic output feeds back into cortex, dG-rabies-GFP was injected into striatal mouth domain CPi.vl.v, and dG-rabies-RFP and PHAL were injected into thalamic mouth domain PF.m. Thalamocortical axons in the MOp-m/i terminate heavily in layer 4 and also in layer 6, overlapping with corticostriatal and corticothalamic neuron dendrites (*inset panels*). C) For unambiguous demonstration that thalamocortical axons feed back onto corticostriatal input neurons in the oro-brachial subnetwork, closing the loop, we implemented channelrhodopsin-assisted circuit mapping. The PF.m was injected with AAV.ChR2 and CPi.vl.v (m/i) was injected with retrobeads. Retrobead-labeled m/i corticostriatal neurons (n=13) were patch clamped and recorded in MOp-m/i during optical stimulation, and the majority (9) responded with an EPSC, while none responded with an IPSC. D) The closed loop schematic.

#### The thalamocortical pathway feeds back into cortical layer 4

The likeliest closed-circuit portion of this subnetwork occurs in the primary motor region MOp-m/i, where thalamocortical axons terminate in the same cortical column as the corticostriatal neurons providing input to the CPi.vl.v (m/i) (Figure 8A). To better visualize the confluence of information from the output of this cortico-basal ganglia subnetwork rising from the thalamus back into the cortex, we injected cell-filling rabies viruses into the CPi.vl.v (m/i) (green glycoprotein-deleted rabies) and PF.m (red glycoprotein-deleted rabies) to label the corticostriatal and corticothalamic neurons, respectively, along with anterograde tracer (PHAL) in the PF.m to see the ascending thalamocortical innervation. The ascending thalamocortical axons terminate densely in layer 4 and moderately in layer 6 of the MOp-m/i (Figure 8B, Supplemental Video 1). The corticothalamic neurons reside primarily in layer 6 with a few scattered in layer 5 (red neurons). The apical dendrites of these corticothalamic neurons extend up to layer 4 where they branch prodigiously (Figure 8B, *insets*) and do not extend above this layer (although the few layer 5 labeled corticothalamic neurons have dendrites that reach layer 1). The corticostriatal neurons primarily populate layer 5 with a few in layer 2/3 (green neurons). They have thick, sparely branched apical dendritic shafts that reach to layer 1 where they branch in finer rami (Figure 8B, *insets*). While these thick apical dendrites pass through the dense field of thalamocortical axon terminals in layer 4, it is unclear from these images if they receive direct synaptic input from this feedback pathway, closing the loop.

#### Cortico-basal ganglia output directly innervates cortico-basal ganglia input neurons in a closed subnetwork loop

To unambiguously determine whether the cortico-basal ganglia subnetwork is truly recurrent, we employed channelrhodopsin-assisted circuit mapping (Petreanu et al. 2007) in the thalamo-corticostriatal segment of the oro-brachial subnetwork. Mice (n=2) were injected with AAV1.hSyn.hChR2.EYFP into the PF.m to label thalamocortical axons with channelrhodopsin and fluorophore-tagged retrobeads into the CPi.vl.v (m/i) to retrogradely label corticostriatal neurons (Figure 8C, *experimental preparation*). Retrobead-labeled cortical neurons were patch-clamped and recorded in an acute slice preparation during blue light stimulation. The bath solution contained tetrodotoxin (1 µM) and 4-aminopyridine (1 mM) to suppress polysynaptic neuronal activity, ensuring only monosynaptic connections could be stimulated and recorded. Recording from layer 5 neurons only (n=13), the majority (9/13) showed an excitatory postsynaptic current to stimulation when clamped at −70 mV (mean±SE, 44.89±14.25 pA; Figure 8C). When clamped at 0 mV, none (0/13) showed an inhibitory postsynaptic current (Figure 8C), as expected since the thalamocortical pathway is a well-known glutamatergic pathway. With 69% of recorded neurons exhibiting a specific monosynaptic excitatory response to thalamocortical stimulation, the cortico-basal ganglia-thalamocortical loop contains a substantial recurrent, closed-circuit component (Figure 8D).

## DISCUSSION

We present for the first time a comprehensive mouse cortico-basal ganglia-thalamic circuit model, with 6 parallel subnetworks (Figure 9A). Previously, we demonstrated that at least fifty-five distinct cortical areas form interconnected cortico-cortical subnetworks: two medial associative subnetworks of exteroceptive visual, auditory, and spatial information, two lateral associative subnetworks of interoceptive limbic areas, and five somatic sensorimotor subnetworks of body regions (Zingg, Hintiryan et al. 2014; Hintiryan, Foster et al. 2016). Each cortico-cortical subnetwork sends largely convergent corticostriatal projections to one (or sometimes several) of ∼30 striatal domains. In this study, we show that the domains of the CP and zones of the ventral striatum project to 14 newly defined SNr domains and 36 GPe domains with complex convergent and divergent projection patterns. Each striatal domain has two efferent streams: the direct pathway to one (or sometimes several) of the nigral domains, with projections from multiple striatal domains converging in a single nigral domain; and the indirect pathway to one or two of the pallidal domains, with far less convergence with other striatal domain outputs. The pallidal domains send projections to the corresponding nigral domain where they re-merge with the matching direct pathway stream from their common striatal source. In turn, the nigral domains project to discrete regions in the PF (and also the VM, Figure 6G-I). Those thalamic output channels project back to cortex, feeding directly back onto corticostriatal neurons in the originating cortical regions in a true closed loop (Figure 8). In a landmark theoretical paper, Alexander and colleagues (1986) postulated multiple parallel networks in monkey, but here we demonstrate them all in one cohesive data set at a much finer resolution using a uniform methodological approach.

**Figure 9.**
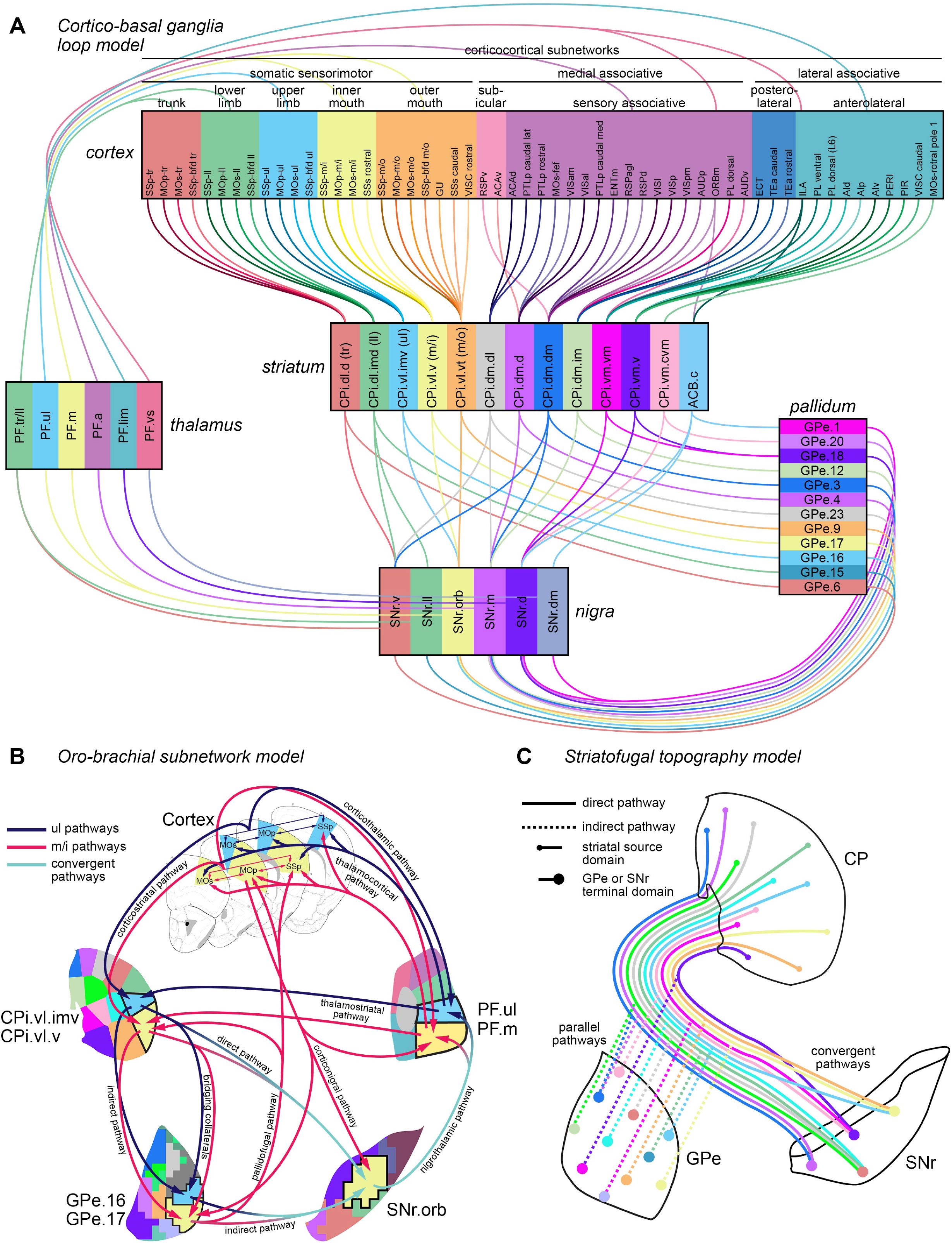
Summary models. A) Cortico-basal ganglia loop model. The corticocortical subnetworks (Zingg, Hintiryan et al. 2014) project into largely distinct striatal subnetworks (Hintiryan, Foster et al. 2016), whose outputs form the parallel indirect (striatopallidal) pathway and the convergent direct (striatonigral) pathway. The pallidal domains send convergent projections to the same nigral domains targeted by their input striatal domains. The nigral domains then project to six regions of the parafascicular thalamus, which in turn are interconnected with the originating cortical regions. Note that (i) only part of the striatum is depicted—the cortical areas have different convergence/divergence patterns in the CPr and CPc domains (ibid.); and (ii) connections between thalamus and cortex may be unidirectional (thalamocortical or corticothalamic) or bidirectional (see Figure 6E for more detail). B) The oro-brachial subnetwork model highlights the precise separation between parallel pathways, the rich interconnectivity within pathways, and the full complexity of the cortico-basal ganglia circuit. C) The striatofugal topography model illustrates the convergence in the direct pathway and parallelism in the indirect pathway. Note that the small and large points represent the true topographic patterns of source and target zones, respectively.

Therefore, our work reveals that the canonical cortico-basal ganglia-thalamic system is composed of a number of parallel subnetworks, each of which is organized by a number of newly identified nodes that are precisely and richly interconnected with each other. A model of the oro-brachial subnetwork (Figure 9B) exemplifies this interconnectvity. This subnetwork integrates upper limb and inner mouth sensorimotor information. The upper limb-specific regions of this subnetwork are interconnected at every node throughout the subnetwork: all of the upper limb cortical areas are heavily connected with each other and all project to the upper limb domain of the CP (Zingg, Hintiryan et al. 2014; Hintiryan, Foster et al. 2016), which in turn projects to the pallidal upper limb domain GPe.16 and the nigral domain SNr.orb; the SNr.orb also receives input from GPe.16 and MOs-ul, and in turn projects to the PF.ul; the PF.ul shares reciprocal connections with the upper limb corticocortical subnetwork and also projects to the upper limb domain of the CP. The inner mouth-specific regions show the same degree of interconnectedness. The upper limb and inner mouth subnetworks are distinctly parallel at every nucleus, except in the corticothalamic projections which show some cross-modality connectivity (Figure 9B, *Cortex*) and in the SNr where the two information streams converge (Figure 9B, *SNr.orb*). Where they merge in the SNr.orb there is the possibility both of neuronal processing of more individual upper limb or inner mouth information, or more integrative information processing as described above (Figure 3D). This may give the SNr the ability to allow the upper limb and inner mouth areas to work separately or coordinate together as need be. Accordingly, in monkey SNr a sample of neurons were recorded during a go/no-go task to correspond to mouth movements, forelimb movements, or both (Schultz, 1986). Our perspective is not intended to deny *any* functional interactivity between distinct subnetworks, as anatomical evidence (including that presented here) has shown crossover between streams of information flow through the basal ganglia (Haber, 2016; Hintiryan, Foster et al. 2016; Aoki et al. 2019). But here we demonstrate that these parallel circuits have the necessary physical structure to operate independently from one another and to coordinate heavily within each subnetwork. This principle applies to the entire cortico-basal ganglia-thalamic loop as shown in this study.

Another remarkable feature we describe is the topographical difference between the direct and indirect pathways. The indirect (striatopallidal) pathway conveys much more of a parallel, point-to-point topography from the CP to the GPe, whereas the direct (striatonigral) pathway exhibits much more convergence of CP efferents (Figure 9C). Note that the points in the diagram in Figure 9C represent the actual topographic patterns of the striatofugal pathways for the source and target domains in the levels depicted. In line with this finding, Kitano and colleagues (1998) recorded neurons in primate SNr that responded to multiple cortical regions with reliable response patterns, such that neurons responsive to the same 2 or 3 particular cortical regions were recorded along much of the rostrocaudal axis of the SNr, suggestive of integration along the length of a nigral domain. On the other hand in GPe, DeLong (1971) recorded from arm- and leg-movement-responsive neurons, and despite finding a partial overlap in their somatotopic representations in the palludim, neuronal responses were associated with either arm or leg movement, never both, indicative of a distinct parallelism. Moreover, in our pallidal analysis, we included a few striatal injections from the intervening region between CPr and CPi; these injections label axons projecting to unique zones of the GPe that no other CP domain projects to, even though these injections have very similar nigral terminal patterns to some of the domains of CPi. Thus we see a highly parallelized striatopallidal topography. While the direct and indirect pathways receive largely similar (though somewhat differently proportioned) cortical inputs (Wall et al. 2013), their output differences suggest differential processing capabilities subserved by the GPe and SNr. However, we noted one important similarity between the two pathways: direct pathway neurons do emit a bridging collateral to the GPe, and this collateral has the same striatopallidal topography as the indirect pathway neurons in the same striatal domain (Figure 5A). The bridging collaterals however have less dense and less extensive terminal fields.

We also revealed direct cortico-nigral projections, which suggest cortical neurons are able to bypass the direct, indirect, and hyperdirect pathways and directly control motor outputs arising from SNr neurons. Several previous anatomical and electrophysiological studies investigated this tract and most failed to find supporting evidence (Rinvik & Walberg, 1969; Goswell & Sedgwick, 1973; Kunzle, 1978; Kitano et al. 1998), although at least one axonal tracing study identified a potential corticofugal terminal field in the SNr (Naito & Kita, 1994). One reason for the mixed findings may be that this direct corticonigral projection pathway appears to be specific to the oro-brachial subnetwork. Its functional significance still remains unclear.

The new pathways we describe here could help to advance investigations of the functions of the basal ganglia. The basal ganglia engages in two main types of reinforcement-related learning, action-outcome and stimulus-response. Action-outcome (A-O) is a goal-directed form of learning wherein an animal associates a behavioral response with a positive or negative consequence; the behavior is performed as a function of the motivational value and obtainability of that outcome (Balleine & O’Doherty, 2010; Smith & Graybiel, 2014). Stimulus-response (S-R) or habit learning is where a reward-associated stimulus can reflexively trigger or motivate a behavioral response that had reliably obtained the reward in the past, even when reward contingency or reward value is degraded (Smith & Graybiel, 2014; O’Hare et al. 2018). Studies in rodents and humans indicate that A-O learning involves the dorsomedial striatum (Yin et al. 2004, 2005; Tanaka et al. 2008), and in rodents it appears that the key region encompasses the domains within CPi.dm and CPc.d (Yin et al. 2005, cf. Hintiryan, Foster et al. 2016). On the other hand, S-R learning involves the dorsolateral striatum (Yin et al. 2004; Tricomi et al. 2009), in what appears to be the domains of the CPi.dl in mouse. While it is known that both A-O and S-R associations are expressed through direct pathway activity (O’Hare et al. 2018), it is not yet known how these two striatal regions interact to resolve behavior. Our findings suggest that one possible locus where direct pathway information streams from both of these striatal regions converge is the SNr.ventral domain. This domain receives convergent inputs from a number of the domains of CPi.dm, CPc.d, and CPi.dl, as well as other somatomotor domains of the CP (Table 1).

Understanding how brain networks change in neuropsychiatric disorders fills in a critical knowledge gap between cellular and molecular pathophysiology on the one end and clinical phenotype on the other, since network dysfunction drives the clinical phenotype (Deisseroth, 2014). A number of neuropsychiatric disorders, such as obsessive-compulsive disorder (Graybiel & Rauch, 2000; Whiteside et al. 2004; Gunaydin & Kreitzer, 2016), major depressive disorder (Nestler et al. 2002), and drug addiction (i.e., Everitt & Robbins, 2016), likely involve alterations in specific subnetworks of the cortico-basal ganglia-thalamic circuit. All together, these studies suggest that a limited number of cortico-basal ganglia-thalamic subnetworks perform specific neurocognitive and neurobehavioral functions, and the specific combination of malfunctions in these circuits underpin complex disorders. Thus a detailed knowledge of the structure and function of these underlying subnetworks is essential.

Finally, a host of neurodegenerative diseases show within-network patterns of degeneration in specific functional and anatomical subnetworks across disease progression (Seeley et al. 2009; Pievani et al. 2011; Fornito et al. 2015). Huntington’s disease (HD) is an example where cortico-basal ganglia-thalamic networks are particularly vulnerable. In HD, an expansion mutation in exon 1 of the Huntingtin (Htt) gene results in a dominant toxic gain-of-function, resulting in motor, cognitive, and psychiatric symptoms that start subtly in midlife and progressively worsen as the neurodegeneration proceeds. Although the striatum is preferentially and profoundly vulnerable to mutant Htt-induced pathology, degeneration and progressive pathological changes are seen in the frontal cortex (Rosas et al. 2006; Harrington et al. 2015), thalamus, including the PF (Heinsen et al. 1999; Kassubek et al. 2005; Tabrizi et al. 2011), and globus pallidus (Rosas et al. 2003, 2006). In later stages of HD, the SNr is degenerated as well (Oyanagi et al. 1989; Vonsattel & DiFiglia, 1998; Douaud et al. 2006; Kiferle et al. 2013). Thus these networked brain regions appear to degenerate in a coordinated, sequential order in HD. Yet there is reason to suspect an even finer pattern of dysfunction and degeneration within the cortico-basal ganglia-thalamic network in HD. Cognitive impairments are among the first measurable signs, appearing years before overt motor signs (Paulsen et al. 2017). Many of the deficits in specific cognitive domains have correlations with unique patterns of morphometric changes in subregions of striatum and cortex, suggesting specific subnetworks are dysfunctional (Harrington et al. 2014). Following cognitive impairment, motor signs then gradually appear and worsen over time, beginning with subtle oculomotor dysfunction and followed by chorea. The chorea can affect the lower limbs, upper limbs, trunk, eye, face, brow, mouth, tongue, and oropharyngeal musculature involved in deglutition (Heemskerk & Roos, 2011; Jankovic & Roos, 2014). As we demonstrated in this study, each of these specific body regions, as well as cognitive function, are regulated by specific subnetworks through the entire cortico-basal ganaglia-thalamo-cortical loop. Dysfunction within each of these subnetworks may prove tractable to experimentation in mouse models of disease. Our new domain maps and subnetwork wiring diagrams provide a structural framework allowing investigation of specific symptom categories, the targeting of interventions to those symptoms, and assessment in alleviating them.

## METHODS

### Subjects and surgeries

Subjects (a total of 249 male 2-month-old wild-type C57Bl6 mice, Jackson Laboratories) were anesthetized with 2% isoflurane in oxygen. Buprenorphine SR (1mg/kg) was administered at the beginning of the surgery as an analgesic. Glass micropipettes (10-30 μm diameter tip) filled with tracer were lowered into the target region and delivered a localized, domain-specific injection (Figure 1D) either by pressure (50 nL volume) or iontophoresis (1-10 min. infusion time, 5 μA, 7 s current pulses). The tracers used were: *Phaseolus vulgaris* leucoagglutinin (PHAL, 2.5%; Vector Laboratories, #L-1110); AAV-GFP (AAV1.hSyn.EGFP.WPRE.bGH, Addgene); AAV-RFP (AAV1.CAG.tdTomato.WPRE.SV40, Addgene); red and green glycoprotein-deleted rabies (Gdel.RV.4tdTomato and Gdel.RV.4eGFP, Wickersham laboratory); AAV1.Syp-GFP.mRuby (Lim laboratory); AAV-Cre (AAV.hSyn.Cre.WPRE.hGH, Addgene); AAVretro-Cre (AAVretro.EF1a.Cre, Salk Institute); Cre-dependent GFP (AAV1.CAG.Flex.eGFP.WPRE.bGH, Addgene); Cre-dependent RFP (AAV1.CAG.Flex.tdTomato.WPRE.bGH, Addgene); fluorogold (FG, 1%; Fluorochrome); and cholera toxin subunit B-AlexaFluor 647 conjugate (CTB, 0.25%; Invitrogen). Animals were monitored daily after surgery until their body weight was on an increasing trajectory. All methods were approved by the Institutional Animal Care and Use Committee of the University of Southern California.

### Roster of injections for striatofugal analyses

All together, 36 anterograde tracer injections were selected and analyzed as representative injections from a data pool of 138 mice with triple anterograde injections (PHAL/AAV-GFP/AAV-RFP) or double coinjections (AAV-GFP/CTB, AAV-RFP/FG), collectively constituting over 300 injections. Three injections were chosen for the nucleus accumbens (ACB), one in the core (ACBc) and two in the shell, medial (ACBsh.m) and lateral (ACBsh.l). The core, medial, and lateral shell have divergent connectivity patterns, and although they mainly connect with a ventral BG network (ventral pallidum, substantia innominata, ventral tegmental area), they also send limited projections to restricted regions of the classical dorsal BG network. We selected injections for 30 domains of the CP, the 29 domains described in Hintiryan, Foster et al. (2016), plus a new subdivision in the caudal extreme (CPext). The CPext previously had a dorsal (CPext.d) and ventral domain, but based on differing input and output patterns, the ventral domain here was split into the rostral ventral (CPext.rv) and caudal ventral (CPext.cv) domains. Additionally, three CP injections were chosen from a level intermediate to CPr and CPi, with projections to regions of the GPe not targeted by any of the other injections included in this experiment. Based on their relative position in the lateral CP, these injection sites appeared to be rostral associations of the somatomotor domains CPi.dl.d (tr), CPi.vl.imv (ul), and CPi.vl.v (m/i); furthermore, their striatonigral projections were highly similar to those three domains (data not shown). Because their striatopallidal projections terminated in regions in the GPe that were targeted by no other CP domains, they were included in the analysis of the GPe data. However, since their projections to the SNr and GPi were homologous to the CPi-level injections’ projections patterns, they were excluded from the analyses of SNr data.

### Histology and imaging

Animal subjects were deeply anesthetized with an overdose bolus of sodium pentobarbital (Euthasol, 2 mg/kg, ip), and cardiac perfused with normal saline followed by 4% boric acid-buffered paraformaldehyde. Brains were post-fixed overnight, embedded in 4% agarose, and sectioned on a vibratome at 50 μm thickness (50-150 μm for rabies-labeled tissues), collected in a 1-in-4 manner into 4 equivalent series, and stored in cryoprotectant at −20°C until staining. Tissue series were stained with rabbit polyclonal anti-PHAL antibody (Vector Labs #AS-2300) at 1:5k and donkey anti-rabbit AlexaFluor 647 (Jackson ImmunoResearch, #711-605-152). Nissl substance was stained with NeuroTrace 435/455 (ThermoFisher, #N21479) at 1:500 to reveal cytoarchitecture. Sections were scanned on an Olympus VS120 epifluorescence microscope with a 10X lens (Plan Apochromat) to capture the Nissl, fluorogold, GFP, RFP, and far red tracers in multichannel photomicrographs; these images were processed for the striatofugal network analysis. High resolution images of some tissue samples (including the rabies labeled tissue from Figures 3D and 8B) were captured with an Andor Dragonfly spinning disk confocal microscope with a 60X lens with a z step of 1 μm.

### Image processing

Captured epifluorescence images were exported as large (14k x 11k pixel) multichannel tiff files (Figure 1E), and subsequently imported into image processing software developed at CIC known as Connection Lens. After an initial atlas matching step, where each section was manually matched to its corresponding level of the Allen Reference Atlas (ARA; Dong, 2007), images containing the pallidum and nigra were registered to the mouse brain atlas (Figure 1F). This work exclusively references levels of the ARA (e.g., ARA 81 refers to Level 81 of the ARA, Bregma=−2.78 mm). Our 1-in-4 series of 50 μm tissue sections gives us a view of the brain that is every-other level in the ARA, itself based on a 1-in-2 series of 100 μm sections. Therefore we registered our tissue sections onto every-other atlas level of the pallidum and nigra, and for this purpose we chose the even levels of the pallidum (i.e., ARA 58-68, even) and odd levels of the nigra (ARA 81-91, odd). All tissue sections were registered to their closest ARA level (i.e., a section containing GPe at ARA 61 was mapped onto either ARA 60 or ARA 62). The determination of which level a given section was assigned to was made by an experienced neuroanatomist. The process of registering to a standardized set of atlas levels provided a uniform dataset that was amenable to computational analysis. After registration, Connection Lens guided users through an interactive segmentation step, creating a binary output image of axons (black) and background (white). Since the images were previously registered, the resulting segmentations were projected onto the atlas frame (Figure 1G). Finally Connection Lens applied an overlap algorithm to quantify the segmented axonal labeling by region (GPe and SNr). Each level of the GPe and SNr in the atlas depicts a single unitary region, yet we knew from the labeling patterns that the striatofugal axonal termination patterns project to a sub-region of each nucleus. We subsequently applied the grid quantification method used previously in our corticostriatal analysis (Hintiryan, Foster et al. 2016), subdividing each nucleus at each atlas level into a square grid space (105 px^2^ per box, equivalent to 63 μm^2^), and quantified the axons per grid box (Figure 1H). Any injection that contributed less than 1500 px to a given level was excluded from the community analysis for that level. A small number of cases had labeling that slightly exceeded the 1500 px threshold for a given level but the labeling was judged too diffuse to be a meaningful terminal field, and were likewise excluded. The surviving grid box data were subjected to network analysis to determine striatonigral and striatopallidal community structure (see next section) (Figure 1I). The derived communities were visualized by recoloring the grid boxes according to community identity (Figure 1J). And finally, projection maps of striatofugal axon terminals were created by aggregating the registered segmented axonal images onto maps of SNr and GPe (Figure 1K).

### Network analysis

The network structure of the dataset was assessed with the Louvain community detection algorithm (Blondel et al. 2008) (Figure 1I), obtained from the Brain Connectivity Toolbox (https://sites.google.com/site/bctnet). Louvain is a greedy, non-deterministic algorithm, with multiple runs producing differing returns of maximized modularity. Importantly, a gamma variable modulates the number of communities detected in a dataset, with smaller gamma values leading to low dimension network structures (fewer nodes, larger communities) and larger gamma values leading to high dimension network structures (more nodes, smaller communities). While the default gamma value of 1 is used commonly (and useful for communicating a frame of reference, as it is a de facto standard), choosing the optimal gamma value is a non-trivial problem.

In an attempt to obtain the most descriptive network partition among this parameterization and variability, we performed a survey of community structures across different gamma values. The Louvain algorithm was run 100 times per each gamma value, over a gamma range of 0 to 2 at 0.05 increments for every nucleus-level. A consensus community structure (conceptually, the “average” community structure; Lancichinetti & Fortunato, 2012) was calculated from each batch of 100 runs at every gamma. The resultant 41 consensus community structures were compiled into a frequency histogram, to determine the most common community structures to arise over the range of gammas for a nucleus-level. The true network structure of the underlying data should act as a ‘magnet’ or attractor for a stable community structure over multiple gammas; therefore an accurately characterized community structure should be obtained across multiple gamma values (Betzel & Bassett, 2017), represented by peaks in the graph (Supplementary Figure 5L,M).

We applied different analytical parameters to the direct (striatonigral) and indirect (striatopallidal) pathway data. The domains in the SNr and GPe exist in 3 dimensions, and for the SNr in particular likely extend across multiple levels of the nucleus. Our goal was to balance across-level similarity with high dimension domain structure, as the input data (i.e., the striatofugal terminal fields) is verified higher dimensional. Moreover, we sought to parse the pallidum and nigra into more than the 3 classically recognized output channels.

For the direct pathway, most of the striatal domains send projections in longitudinal columns along the entire rostrocaudal extent of the SNr. This suggests there is a relatively high degree of consistency in the community structures across adjacent levels of the SNr. However, caudally the SNr becomes physically smaller in cross-sectional area and the axonal terminal fields exhibit a higher degree of convergence than in rostral levels (Supplementary Figure 2A, 5B,C; also see Hedreen & DeLong, 1991). We quantitatively characterized this convergence to describe the degree of integration, and utilized these data to inform our selection of gamma values. Using the quantified grid box data, we created frequency distributions of the boxes categorized by how many different striatal inputs they received (only boxes receiving input were included in this analysis) (Supplementary Figure 5A). We applied a minimum threshold such that inputs contributing less than 5% overlap to a box (551 px in a 105 px^2^ box) were excluded from that box’s tally of inputs. When graphed together, the histograms show that the rostral SNr levels 81-85 have negatively skewed distributions, while the caudal levels 87-91 have more platykurtic distributions with relatively fatter tails in the 10-15 inputs per box range (Supplementary Figure 5B), a trait that becomes more apparent when the rostral levels and caudal levels are averaged (Supplementary Figure 5C). There is no significant correlation of mode (peak value) of each histogram with rostrocaudal level as assessed by linear regression (r=.4252; P=.4006; Supplementary Figure 5E), and there is no significant difference in mode between the rostral (mean±SE: 7.0±.577) and caudal (7.3±1.856) groups (P=.8796; t=.1715; df=4; 2-tailed Welch’s t test), verifying that the graphs have similar central tendencies (Supplementary Figure 5C). Two-way ANOVA of the rostral and caudal groups (SNr level x amount of integration) finds a significant main effect of integration (P<.0001; df=14; F=12.46) and a significant interaction of integration and SNr level (P=.0219; df=14; F=2.136). Post hoc Bonferroni tests reveal the caudal group has a significantly greater proportion of boxes receiving 11 inputs (P<.0033; t=3.385) and a nearly significant difference for 10-input boxes (.05>P>.025, familywise-adjusted α=.0033; t=2.278). Thus the caudal 3 levels of the SNr have a significantly greater proportion of their area devoted to high integration (10 or more inputs per box) (Supplementary Figure 5D,F), indicating that the rostral and caudal SNr should be analyzed with different parameters since there is likely a different, more integrative domain structure caudally.

Since the integration analysis indicates that the rostral and caudal halves of the SNr form two groups, we selected community structures that were most common through the rostral and caudal groups (Supplementary Figure 5L). By stacking the histograms of the component levels, the peaks in the graphs reveal the most stable community structures common to all constituent levels. For the rostral SNr group, there were two clear peaks, for a 6-domain and a 10-domain structure (Supplementary Figure 5L, *SNr rostral group*). We have selected to present the 10-domain structure, because we sought to subdivide the SNr into as fine a coherent partition as possible, although the 6-d structure may also be a valid way to interpret the striatonigral data as well, since it is possible there is a nested multi-scale network architecture to the striatonigral pathway as with the corticostriatal pathway. For the caudal group, there was a clear peak for the 7-domain structure (Supplementary Figure 5L, *SNr caudal group*). All gamma values returning the chosen domain structure for each nucleus-level were pooled and the consensus community structure of that pool was the final network output. After determining the community structure for each level of the SNr, we joined together communities on adjacent levels based on continuity and similarity of inputs (Table 1).

Each indirect pathway input tends to densely innervate just a few levels of the GPe. GPe boxes integrate at most 9 inputs, much more restricted than the SNr (up to 15), and mean mode of the GPe (3.33±.558 [mean±SE]; n=6) is significantly smaller than mean mode of SNr (7.17±.872; n=6), indicating less integration and more segregated relaying of striatal activity through the indirect pathway (P=.0060; t=3.702; df=8; 2-tailed Welch’s t test). The frequency distributions of inputs per box show similarly shaped histograms across GPe levels, with decreasing mode value from rostral to caudal (Supplementary Figure 5G,H). Linear regression of the histogram modes shows a significant correlation with GPe level (r=.8607; P=.0278), with mode decreasing towards caudal levels (Supplementary Figure 5J), indicating that caudal GPe integrates progressively fewer inputs per box (Supplementary Figure 5I,K). Given that the graphs vary continuously along the rostrocaudal axis, and the lower degree of continuity of striatopallidal projections along that axis, we evaluated each level of the GPe independently (Supplementary Figure 5M).

### Whole brain 3D imaging and reconstruction of neuronal morphology

Intact brains were SHIELD-cleared as described by Park and colleagues (2019), placed in refractive index matching solution (EasyIndex, LifeCanvas), and imaged on a LifeCanvas lightsheet microscope at 4X and 10X magnification. These images as well as z stack images captured with the DragonFly confocal microscope were viewed within Aivia reconstruction software (v8.8.2. DRVision) and neurons were manually reconstructed. Geometric processing of the reconstructions was performed using the Quantitative Imaging Toolkit (http://cabeen.io/qitwiki), and morphometric data were obtained from the reconstructions with NeuTube (v1.0z). Descriptive statistics of the morphological features of these neurons were generated by NeuTube.

### Channelrhodopsin-assisted circuit mapping

Wild-type mice (n=2) were injected with AAV expressing channelrhodopsin (pAAV.hSyn.hChR2(H134R).EYFP, Addgene 26973, titer 2.5e13) into the PF thalamic nucleus, and rhodamine-conjugated retrobeads (Lumfluor) into the CPi.vl.v (m/i). Three weeks following the injections, acute brain slices containing MOp were prepared for recording. Following anesthesia, the animal was decapitated, and the brain was quickly removed and immersed in ice-cold dissection buffer (composition: 60 mM NaCl, 3 mM KCl, 1.25 mM NaH2PO4, 25 mM NaHCO3, 115 mM sucrose, 10 mM glucose, 7 mM MgCl2, 0.5 mM CaCl2; saturated with 95% O_2_ and 5% CO_2_; pH = 7.4). Brain slices of 300 μm thickness containing the MOp were cut in a coronal plane using a vibrating microtome (Leica VT1000s). Slices were allowed to recover for 30 min in a submersion chamber filled with warmed (35°C) ACSF and then cooled gradually to room temperature until recording. The presence of retrobead (RFP) and hChR2 (YFP) labeling in the frontal cortex was examined under green and blue fluorescence bulbs in the corresponding slices before recording. Red-labeled MOp layer V neurons were visualized under a fluorescence microscope (Olympus BX51 WI). Patch pipettes (Kimax) with ∼6-7 MΩ impedance were used for whole-cell recordings. Recording pipettes contained: 130 mM K-gluconate, 4 mM KCl, 2 mM NaCl, 10 mM HEPES, 0.2 mM EGTA, 4 mM ATP, 0.3 mM GTP, and 14 mM phosphocreatine (pH, 7.25; 290 mOsm). Signals were recorded with a MultiClamp 700B amplifier (Molecular Devices) under voltage clamp mode at a holding voltage of –70 mV for excitatory currents and 0 mV for inhibitory currents, filtered at 2 kHz and sampled at 10 kHz. 1 μM tetrodotoxin (TTX) and 1 mM 4-aminopyridine (4-AP) was added to the external solution for isolation and recording of monosynaptic responses to blue light stimulation (5 ms pulse, 3 mW power, 5 trials, delivered via a mercury Arc lamp gated with an electronic shutter).

### Anterograde transsynaptic tracing

Detailed methodology can be found in Zingg and colleages (2017, 2020). In brief, anesthetized mice were iontophoretically injected with Cre-dependent AAV-tdTomato (pAAV.CAG.Flex.tdTomato.WPRE, Addgene) in the rostral medial SNr, and then pressure injected (50 nL) with AAV-Cre (AAV.hSyn.Cre.WPRE.hGH, Addgene) in the medial ACB. The AAV-Cre is transported anterogradely down the axons and is released from the terminals, where it infects postsynaptic cells that have been infected with high concentrations of Cre-dependent AAV-tdTomato, unlocking strong fluorophore expression in those downstream neurons. After a 3-week post-operative recovery, animals were pentobarbital-anesthetized and perfused as above. The Cre injection site was verified by staining with mouse anti-Cre recombinase monoclonal antibody (Millipore Sigma, MAB3120) and donkey anti-mouse AlexaFluor 647 (Jackson ImmunoResearch, #715-605-150).

### Fluorescence micro-optical sectioning tomography (fMOST)

All fMOST experiments were conducted in accordance with the Institutional Animal Ethics Committee of Huazhong University of Science and Technology. For sparse labeling of striatal neurons, we created a single pAAV co-package of DNA cassettes of CMV.Cre and Cre-dependent EF1a.DIO.GFP at a ratio of 1:1,000,000, respectively, so that the final viral admixture contained one virus with both cassettes for every 1,000,000 viruses that contained only the EF1a.DIO.GFP cassette (total viral concentration 8e12 gc/ml, from BrainVTA Co., Ltd., Wuhan, China). This admixture was pressure injected (100 nL) into CPi.dm (M-L, A-P, D-V: 0.14,-1.3,-2.6), allowing high viral load for the fluorophore gene with low frequency of co-expression of Cre, resulting in sparse yet bright GFP expression. A detailed protocol has been previously described (Sun, et al., 2019). After 5 weeks, mice were anesthetized, perfused with 0.01M PBS and 4% paraformaldehyde, and brains were post-fixed overnight. For whole brain imaging, brains were rinsed in 0.01M PBS solution and dehydrated in a graded ethanol series (50, 70 and 95% ethanol), submerged in gradient series of Lowicryl HM20 resin, and polymerized at a gradient temperature in a vacuum oven.

The resin-embedded whole brain samples were imaged using fMOST, a three-dimensional dual-wavelength microscope-microtome combination instrument (see Gong et al. 2016 for a detailed description). Block imaging mode was used to slice and scan layer by layer through the whole sample in the coronal plane. GFP-labeled neurons and propidium iodide-stained cytoarchitecture were acquired at a voxel resolution of 0.32×0.32×1 μm^3^. The raw images were first preprocessed for intensity correction, and then the image sequence was converted to TDat, an efficient 3D file format for large volume images, to facilitate the computing of terabyte- and petabyte-scale brain-wide datasets (Li et al. 2017). We employed GTree for semi-automated, manually-assisted reconstruction of neuronal morphology in 3D (Zhou et al. 2018). Subsequently, neuronal morphological data were mapped to the Allen CCFv3 using BrainsMapi, a robust image registration interface for large volume brain images (Ni et al. 2020). Because the contours of brain regions on the propidium iodide-stained images can be more clearly identified, this greatly reduces the difficulty of accurate registration.

### D1 and A2A cell type specific tracing

For labeling D1 and D2 dopamine receptor-expressing medium spiny neurons (MSNs), adult Adora2a-cre (GENSAT 036158-UCD, for labeling D2-MSNs) and Drd1a-cre (GENSAT 017264-UCD, for D1-MSNs) congenic mice on a C57BL6/JL background were obtained from GENSAT and backcrossed with wild type C57BL mice (Jackson Laboratories) for several generations. Surgeries were performed between 10-12 weeks of age. Subjects were anesthetized with 2% isoflurane in oxygen. Buprenorphine SR (1mg/kg) was administered at the beginning of the surgery as an analgesic. Glass micropipettes (10-30 µm diameter tip) filled with AAV-DIO-EGFP (AAV-DJ.hSyn-DIO-EGFP.WPRE.bGH, Lim lab) were lowered into the target region and delivered a localized injection by pressure (50-150 nL volume) at a rate of 100 nL/min. After 3 weeks, animals were deeply anesthetized and perfused. Tissue sections were stained with DAPI and imaged on an Olympus VS120 epifluorescence microscope with a 10X objective lens. All procedures to maintain and use mice were approved by the Institutional Animal Care and Use Committee (IACUC) at the University of California, San Diego.

### Code availability

The code for Connection Lens software is not currently available due to pending preparation of a manuscript describing it in detail.

## Acknowledgements

The authors wish to thank technical support from Stanley Kong, Kendall Kirio, Jennifer Gonzalez, Fernando Sanchez Cano, Zachary Hobel. Funding for this research provided by: NIH BICCN Grant U01 MH114829 (HWD), R01 MH094360-06 (HWD), NIH U19MH114821, U19MH114831; The National Science Fund for Creative Research Group of China (grants 61721092, 61890953, and 61890954).

---

**Supplementary Figure 1.**
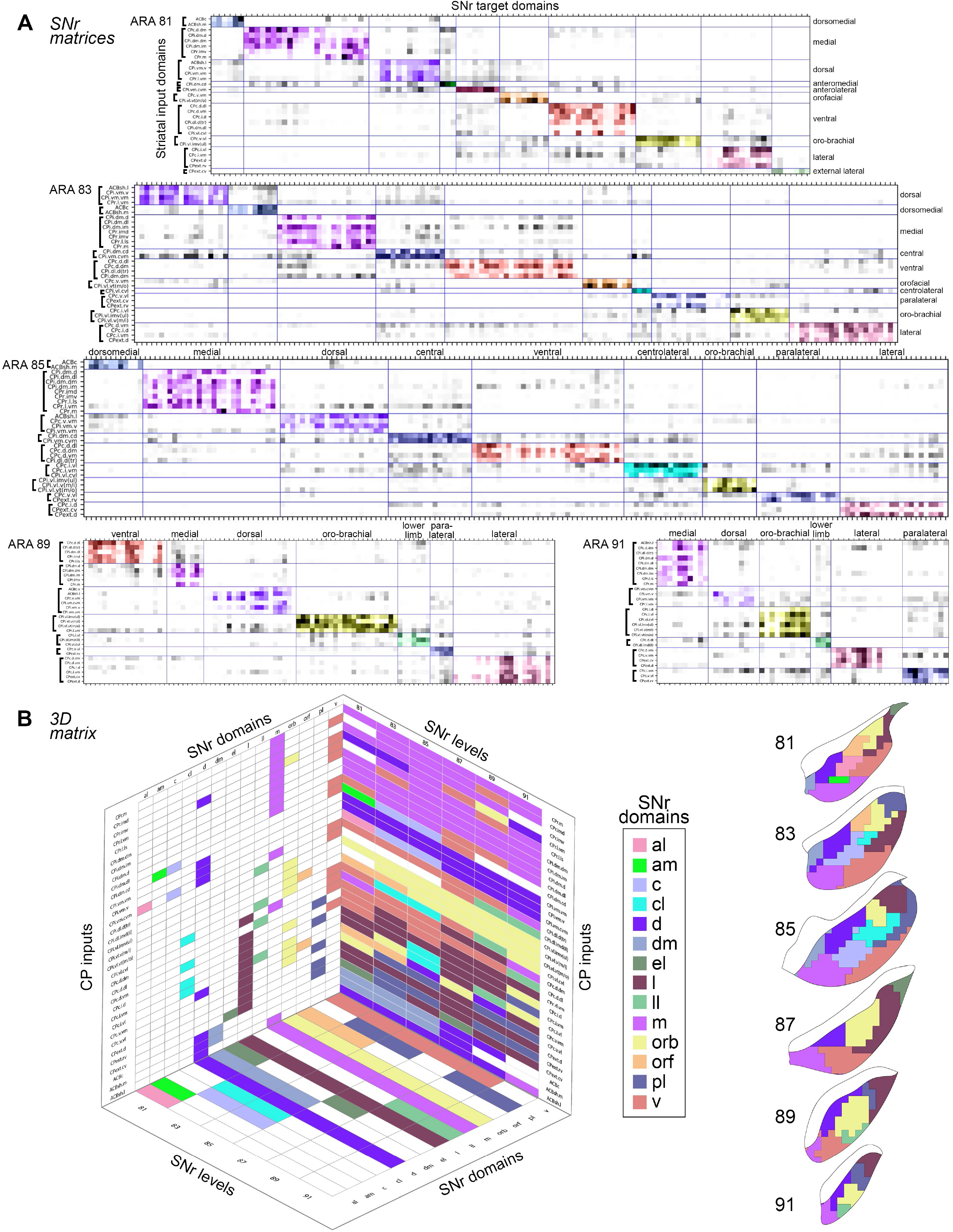
SNr matrices. A) Ordered matrices depict the community structure determined by the Louvain algorithm. The striatal input domains (rows) terminate in a common set of nigral grid boxes (columns) at each nucleus-level. The new nigral domains lie along the diagonal and are colored to match the SNr domain map (shown below in (B), *right*). The matrix for SNr 87 is shown in Figure 2G. B) The 3D matrix shows the striatal input domains on the vertical axis, identifying to which SNr domains they contribute (*lefthand plane*); and to which level of the SNr they provide input (*righthand plane*), color-coded to indicate the domain they contribute to, with white cells indicating no contribution at that level. Each SNr domain is also plotted according to which SNr levels it is present at (*bottom plane*). The SNr domain map is presented at right for reference.

**Supplementary Figure 2.**
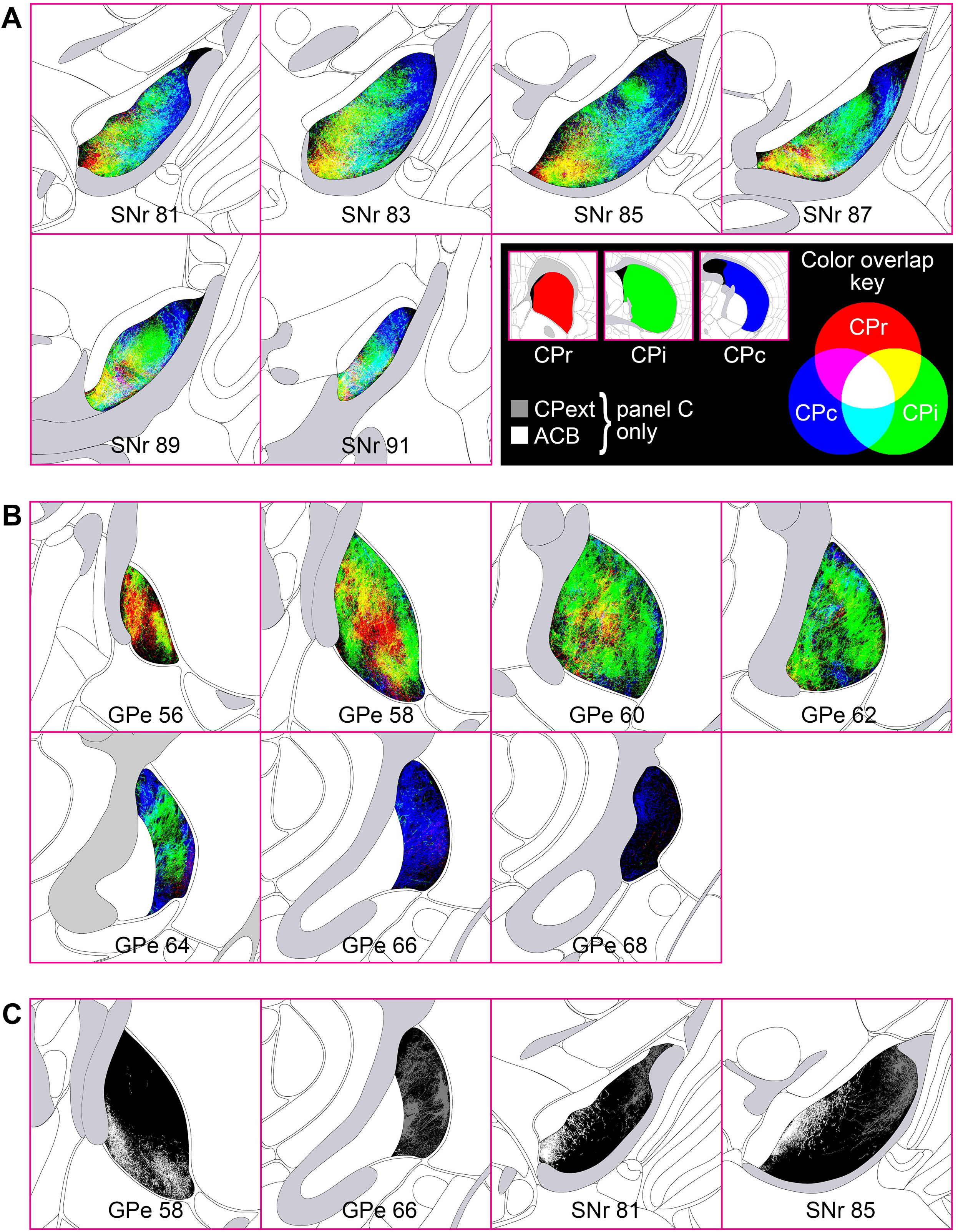
Division-level striatofugal projection maps, blended-color style. Similar to the division-level maps presented in Figures 2A and 4A, the maps here present the division-level termination patterns, but when axons from different divisions overlap, the pixel color is a blended RGB value of the constituent axons (c*olor overlap key*). The maps in Figures 2A and 4A were made with opaque layers (CPr>CPi>CPc, top to bottom) such that pixels of axon in the upper layers occlude any other axons in lower layers in the same pixel; those maps were meant to emphasize more of the differences in division-level terminal patterns, whereas the maps in this figure emphasize the areas of overlap within SNr (A) and GPe (B). The terminations of the ACB and CPext (also known as the tail of the caudate) are shown in grayscale in (C).

**Supplementary Figure 3.**
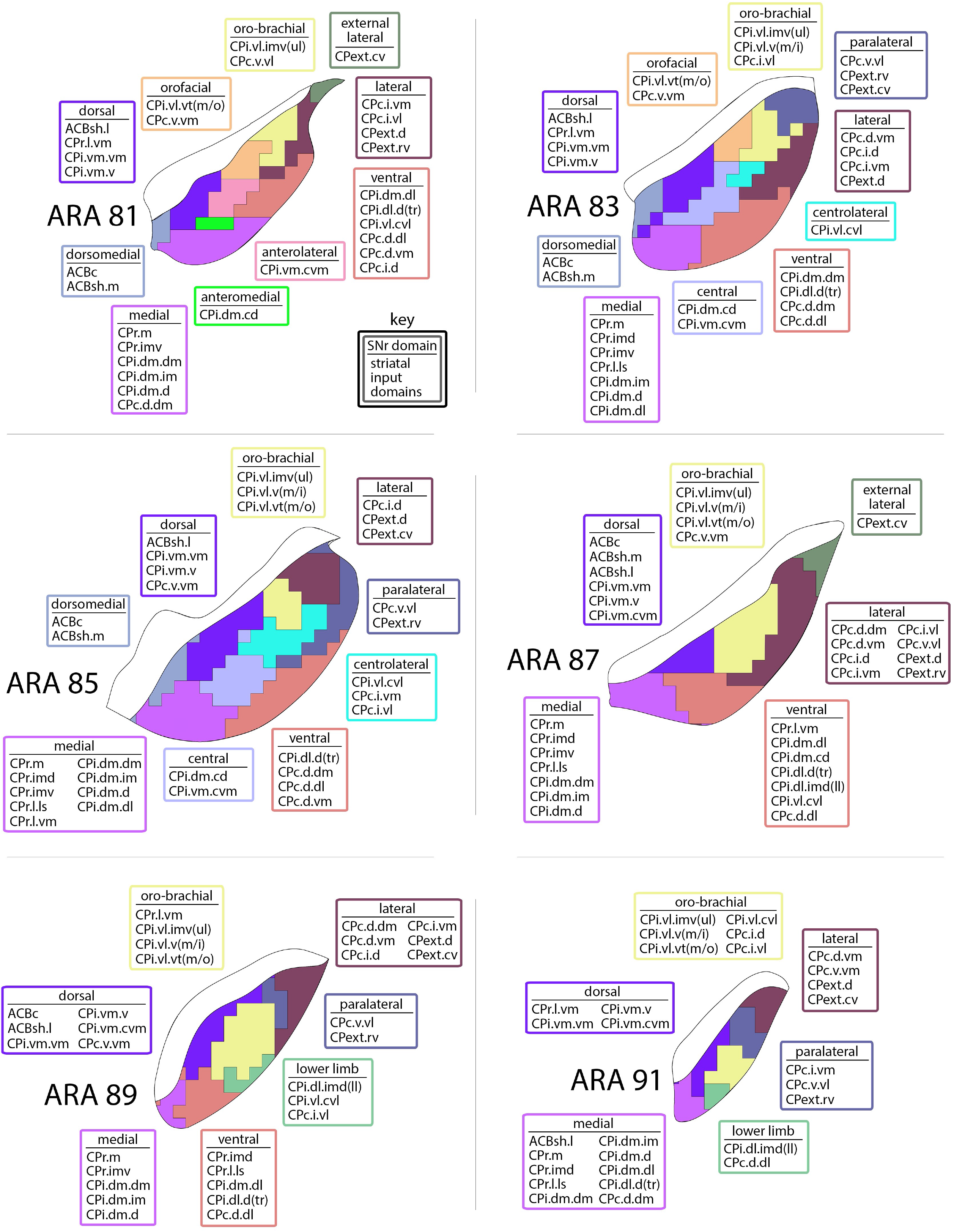
SNr input map. At each level of the SNr, the striatal inputs to each nigral domain are listed in the boxes.

**Supplementary Figure 4.**
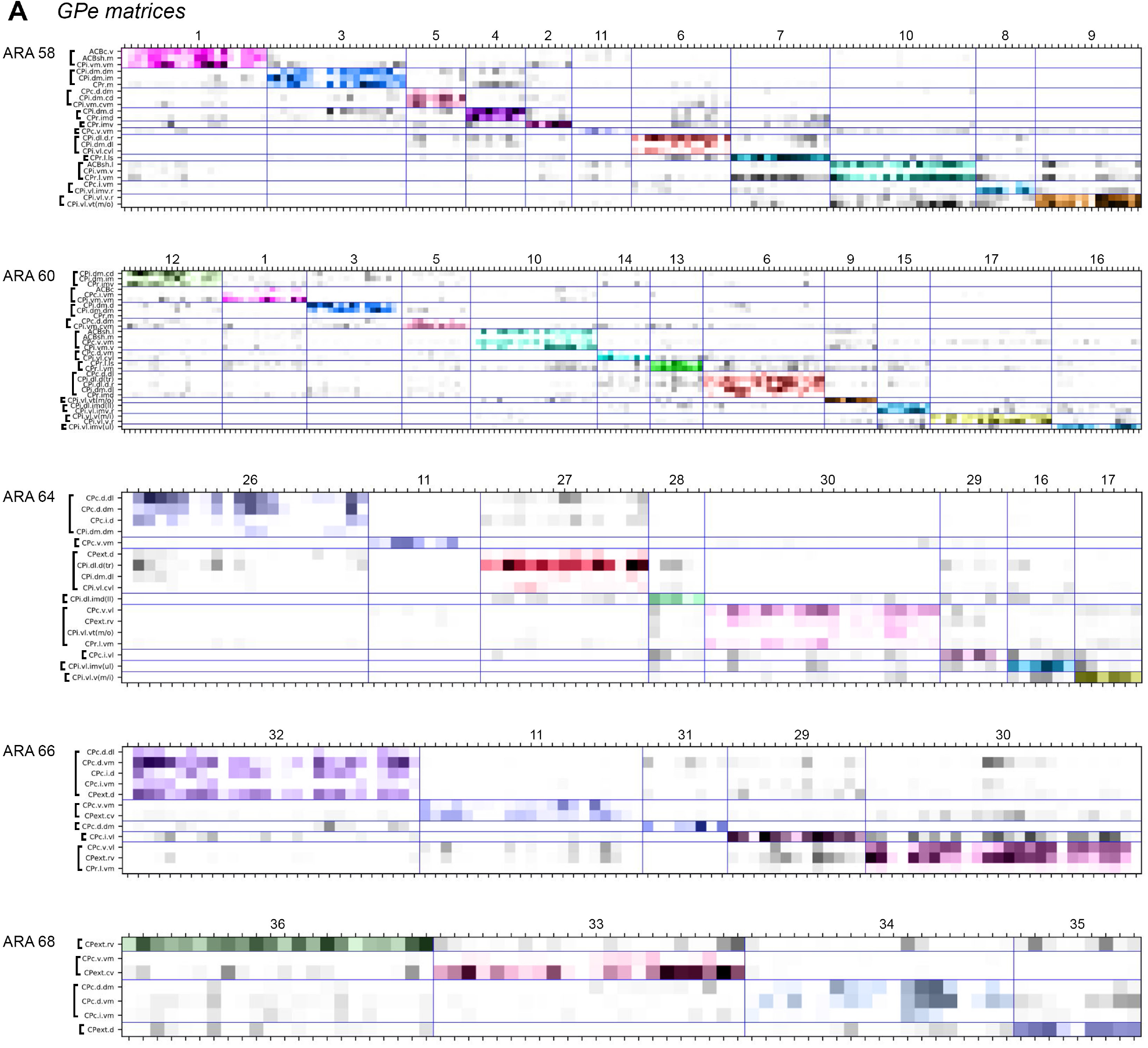
GPe matrices. A) Ordered matrices depicting the community structure determined by the Louvain algorithm. The striatal input domains (rows) terminate in a common set of pallidal grid boxes (columns) at each nucleus-level. The new pallidal domains lie along the diagonal and are colored to match the GPe domain map (shown in Figure 4F). The domain numbers are listed along the top. The matrix for GPe 62 is shown in Figure 4E.

**Supplementary Figure 5.**
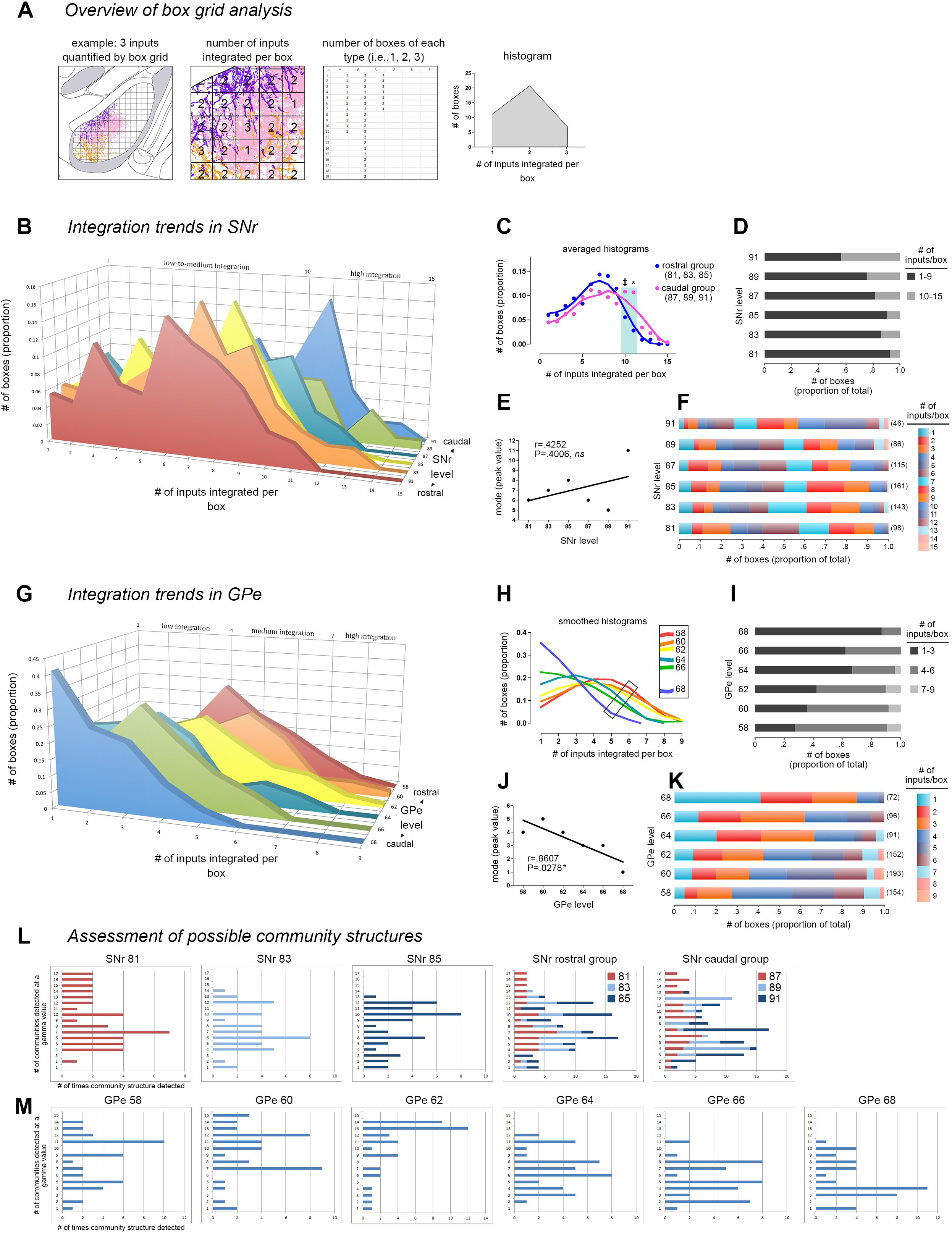
Box grid data describes integration trends in the pallidum and nigra and informs the community analysis. The example in (A) illustrates how the box grid analysis works using 3 inputs (pink, purple, and orange; *left panel*). Each whole and partial box in the ROI receives 1-3 inputs (*center panel*); the number of boxes of each input category is tallied (*right panel*) and plotted in a frequency distribution (*histogram*). B) Frequency distributions for each level of the SNr are shown rostral to caudal (front to back in the graph). The caudal 3 levels have a greater proportion of boxes devoted to integrating high numbers of inputs, as seen by the tails of their distributions sticking out in that range. This was validated statistically by comparing the caudal 3 and the rostral 3 distributions with ANOVA (* P<0.0033, ‡ 0.05>P>0.025), which are averaged and summarized in (C). The proportion of boxes integrating various numbers of inputs at each level is represented in the stacked bar charts: (D) shows a categorized graph, with low-to-moderate integration (1-9 inputs) in black and high integration (10-15 inputs) in gray; (F) shows the same data with each bin in a different color (the bars from left to right correspond to the legend from top to bottom), and the numbers in parentheses at the end of each bar are the total number of boxes at each level. E) Regression analysis of the peak of each histogram from (B) with SNr level shows no correlation (r=0.4252, P=0.4006). G) Frequency distributions for each level of GPe are shown caudal to rostral (front to back in the graph). The distributions are fairly similar in shape and exhibit a linear trend, from rostral GPe integrating higher numbers of inputs per box and shifting to successively lower integration with each caudal level. Smoothing of the histograms and plotting them in a single plane in (H) highlights this stepwise trend from higher to lower integration, rostral to caudal, respectively (*inset*). J) This trend was found to be significant when assessed by regression analysis (r=0.8607, P=0.0278). Stacked bar graphs in (I) and (K) are as in (D) and (F). L,M) The Louvain community detection algorithm was run multiple times over a range of gamma values, with gamma modulating the number of communities (i.e., domains) detected in the nigra or pallidum. The bar graphs show the different community structures detected and how many times they were detected; each integral increment on the x axis means one gamma detected that community structure, so the peaks represent the most commonly detected community structures. These survey analyses were run for each nucleus-level, and results for SNr 81-85 can be seen individually and stacked together (*SNr rostral group*), along with the stacked graph for the SNr caudal group (SNr 87-91, individual graphs not shown). The individual graphs for GPe are shown in (M).

**Supplementary Figure 6.**
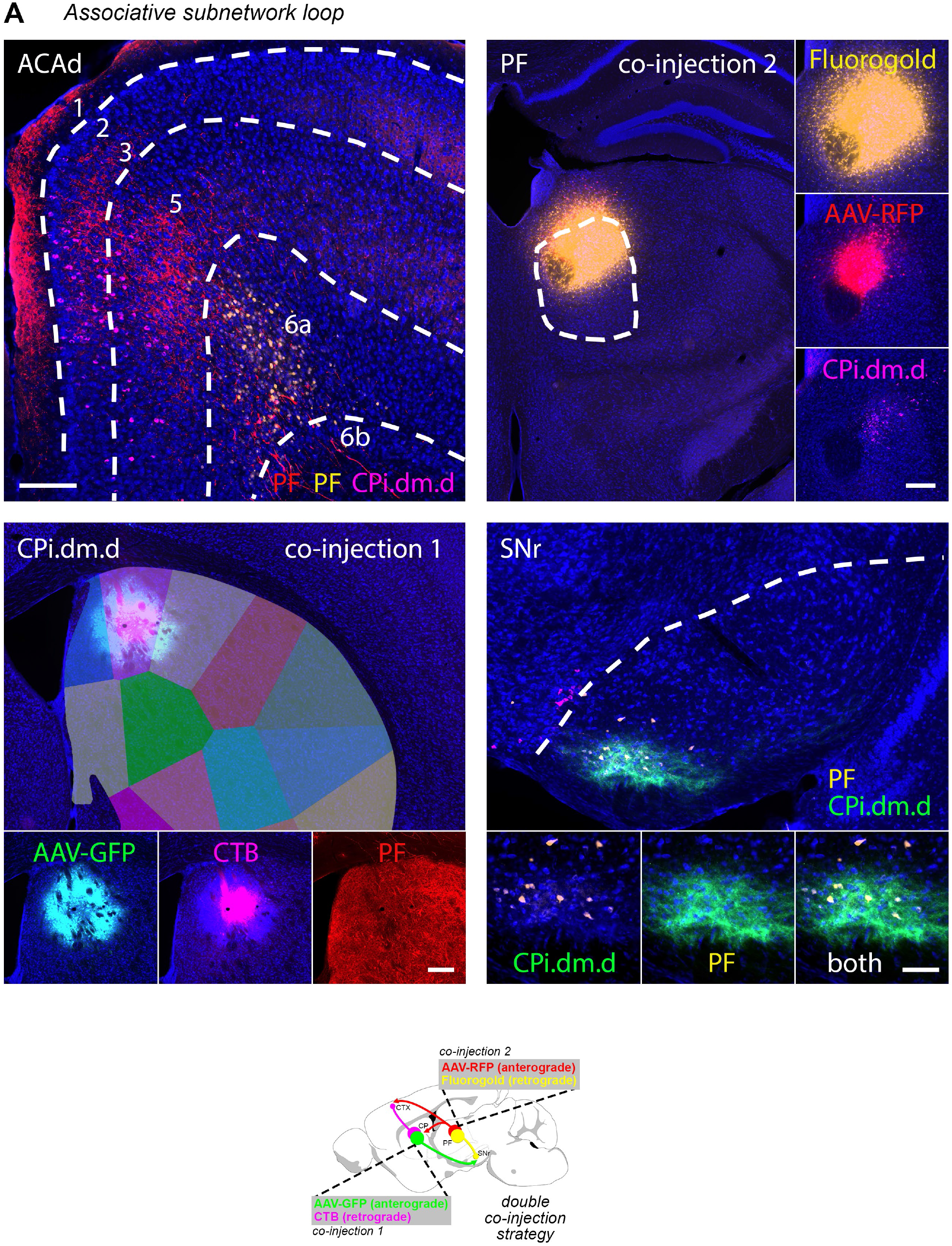
Double co-injection demonstrating the associative subnetwork. A) One co-injection was placed in the striatum (CPi.dm.d, injected with AAV-GFP and CTB) and the other co-injection was made in the thalamus (PF.a, injected with AAV-RFP and fluorogold). The transported tracers overlap in the cortex (ACAd) and nigra (SNr.m), revealing the discrete, closed-loop nature of the subnetwork. Note that the RFP-labeled axons in the striatum and CTB-labeled somata in the PF indicate the direct thalamostriatal pathway between those two nodes. The injection strategy and prototypical circuit schematic are shown in (B).

**Supplementary Figure 7.**
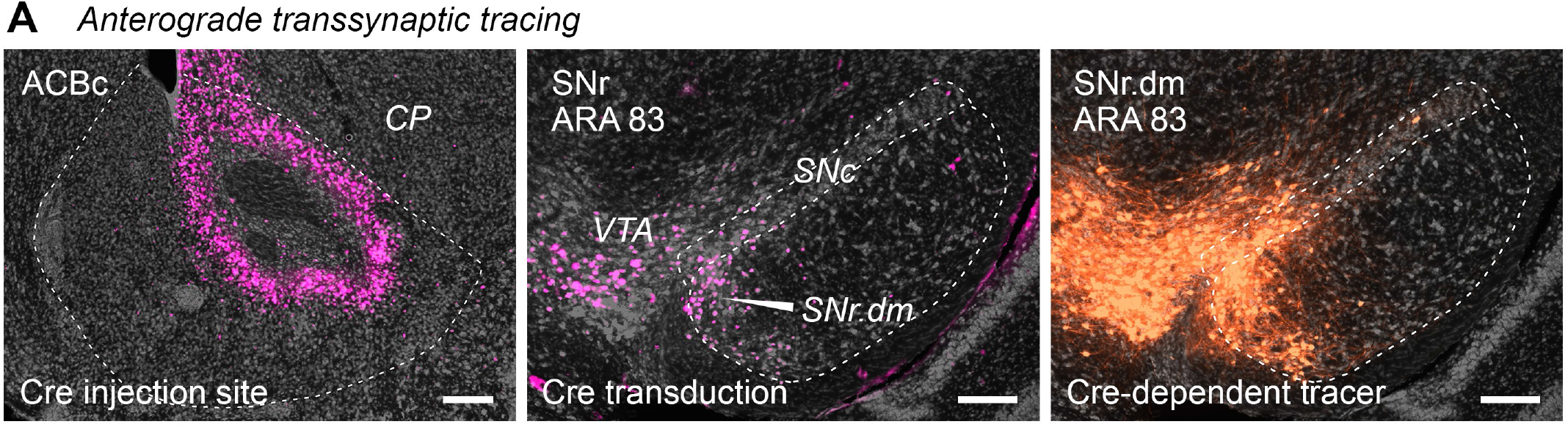
Anterograde transsynaptic tracing. A) AAV.Cre was injected into the nucleus accumbens core (*left panel*). Cre is visualized with anti-Cre recombinase antibody (pink labeling). Anterograde transsynaptic transduction of neurons can be seen in SNr.dorsomedial (*center panel*). Cre expression in neurons in substantia nigra pars compacta (SNc) and ventral tegmental area (VTA) may be due to either anterograde transsynaptic transduction or retrograde transduction, since both areas are reciprocally connected and AAV can infect retrogradely at low efficiency (unpublished observation). But the ACB to SNr pathway is unidirectional, and so this labeling is due to anterograde transmission. Cre-dependent fluorophore expression (orange labeling) is seen in the transduced Cre-expressing neurons of the SNr, SNc, and VTA (*right panel*). Only the SNr labeling accounts for the axons labeled in PF.vs and VM.vs (Figure 6D,G, *images labeled ‘SNr.dm’*), because the catecholaminergic neurons of SNc and VTA do not project to either thalamic nucleus. Scalebar = 200 µm.

**Supplementary Figure 8.**
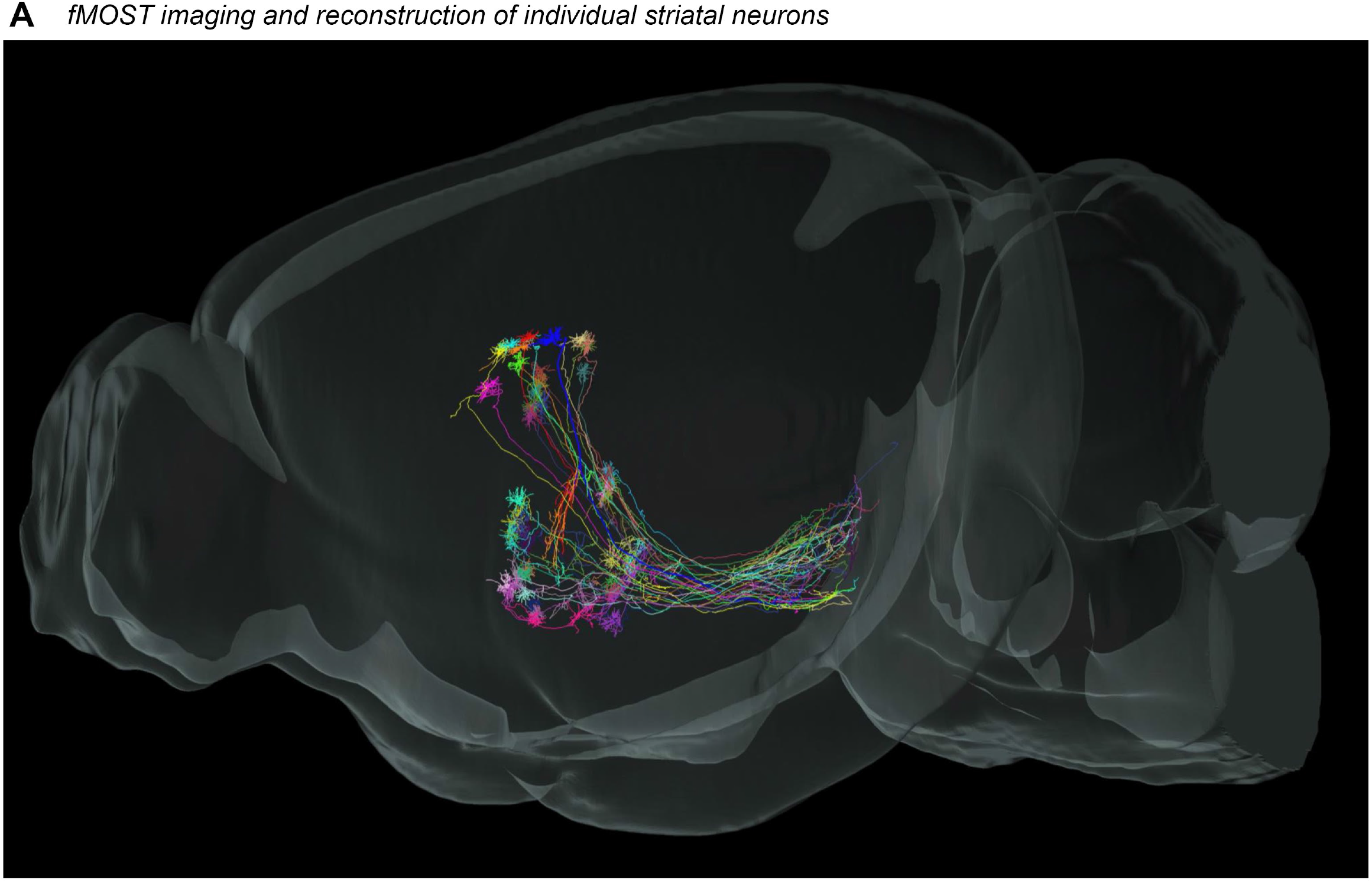
fMOST data. A) GFP expression was sparsely induced in 30 striatal neurons, imaged with the fMOST system, reconstructed, and registered to the Allen Common Coordinate Framework version 3. This sagittal view of the data shows striatofugal axons in pallidum and nigra.

## REFERENCES

Abdi, A, Mallet, N, Mohamed, FY, Sharott, A, Dodson, PD, Nakamura, KC, Suri, S, Avery, SV, Larvin, JT, Garas, FN, Garas, SN, Vinciati, F, Morin, S, Bezard, E, Baufreton, J, Magill, PJ (2015) Prototypic and arkypallidal neurons in the dopamine-intact external globus pallidus. Journal of Neuroscience, 35(17):6667–6688

Alexander, GE, DeLong, MR, Strick, PL (1986) Parallel organization of functionally segregated circuits linking basal ganglia and cortex. Annual Review of Neuroscience, 9:357–381

Alexander, GE, Crutcher, MD, DeLong, MR (1990) Basal ganglia-thalamocortical circuits: Parallel substrates for motor, oculomotor, “prefrontal” and “limbic” functions. Progress in Brain Research, 85:119–146

Aoki, S, Smith, JB, Li, H, Yan, X, Igarashi, M, Coulon, P, Wickens, JR, Ruigrok, TJH, Jin, X (2019) An open cortico-basal ganglia loop allows limbic control over motor output via the nigrothalamic pathway. eLife, 8:e49995

Bakhurin, KI, Goudar, V, Shobe, JL, Claar, LD, Buonomano, DV, Masmanidis, SC (2017) Differential encoding of time by prefrontal and striatal network dynamics. Journal of Neuroscience, 37(4):854–870

Balleine, BW, O’Doherty, JP (2010) Human and rodent homologies in action control: corticostriatal determinants of goal-directed and habitual action. Neuropsychopharmacology Reviews, 35:48–69

Betzel, RF, Bassett, DS (2017) Multi-scale brain networks. NeuroImage, 160:73–83

Blondel, VD, Guillaume, J-L, Lambiotte, R, Lefebvre, E (2008) Fast unfolding of communities in large networks. Journal of Statistical Mechanics: Theory and Experiment, 10:1–12

Cazorla, M, Delmondes de Carvalho, F, Chohan, MO, Shegda, M, Chuhma, N, Rayport, S, Ahmari, SE, Moore, H, Kellendonk, C (2014) Dopamine D2 receptors regulate the anatomical and functional balance of basal ganglia circuitry. Neuron, 81:153–164

Cui, G, Jun, SB, Jin, X, Pham, MD, Vogel, SS, Lovinger, DM, Costa, RM (2013) Concurrent activation of striatal direct and indirect pathways during action initiation. Nature, 494:238–242

Deisseroth, K (2014) Circuit dynamics of adaptive and maladaptive behavior. Nature, 505:309–317

DeLong, MR (1971) Activity of pallidal neurons during movement. J. Neurophysiol. 34(3):414–427

DeLong, MR, Strick, PL (1974) Relation of basal ganglia, cerebellum, and motor cortex units to ramp and ballistic limb movements. Brain Research, 71(2-3):327–335

Deniau, JM, Chevalier, G (1985) Disinhibition as a basic process in the expression of striatal functions. II. The striato-nigral influence on thalamocortical cells of the ventromedial thalamic nucleus. Brain Research, 334:227–233

Deniau, JM, Feger, J, Le Guyader, C (1976) Striatal evoked inhibition of identified nigro-thalamic neurons. Brain Research, 104(1):152–156

Ding, L, Gold, JL (2013) The basal ganglia’s contributions to perceptual decision making. Neuron, 79:640–649

Dirnberger, G, Jahanshahi, M (2013) Executive dysfunction in Parkinson’s disease: a review. Journal of Neuropsychology, 7:193–224

Dong, HW (2007) The Allen Reference Atlas: a digital color brain atlas of the C57Bl/6J male mouse. Wiley

Douaud, G, Gaura, V, Ribeiro, M-J, Lethimonnier, F, Maroy, R, Verny, C, Krystkowiak, P, Damier, P, Bachoud-Levi, A-C, Hantraye, P, Remy, P (2006) Distribution of grey matter atrophy in Huntington’s disease patients: a combined ROI-based and voxel-based morphometric study. NeuroImage, 32(4):1562–1575

Everitt, BJ, Robbins, TW (2016) Drug addiction: updating actions to habits to compulsions ten years on. Annual Review of Psychology, 67:23–50

Flaherty, AW, Graybiel, AM (1993) Output architecture of the primate putamen. J Neurosci, 13(8):3222–3237

Fornito, A, Zalesky, A, Breakspear, M (2015) The connectomics of brain disorders. Nature Reviews Neuroscience, 16:159–172

Fujiyama, F, Sohn, J, Nakano, T, Furuta, T, Nakamura, KC, Matsuda, W, Kaneko, T (2011) Exclusive and common targets of neostriatofugal projections of rat striosome neurons: a single neuron-tracing study using a viral vector. European J Neurosci, 33:668–677

Gerfen, CR (1984) The neostriatal mosaic: compartmentalization of corticostriatal input and striatonigral output systems. Nature, 311:461–464

Gerfen, CR (1985) The neostriatal mosaic. I. Compartmental organization of projections from the striatum to the substantia nigra in the rat. J Comp Neurol, 236:454–476

Gimenez-Amaya, JM, Graybiel, AM (1990) Compartmental origins of the striatopallidal projection in the primate. Neuroscience, 34(1):111–126

Gittis, AH, Kreitzer, AC (2012) Striatal microcircuitry and movement disorders. Trends in Neurosciences, 35(9):557–564

Gong, H, Xu, D, Yuan, J, et al. (2016) High-throughput dual-colour precision imaging for brain-wide connectome with cytoarchitectonic landmarks at the cellular level. Nature Communications, 7:12142

Goswell, MJ, Sedgwick, EM (1973) Is there a cortico-nigral tract? Some neurophysiological observations. Brain Research, 50:437–440

Graybiel, AM, Rauch, SL (2000) Toward a neurobiology of obsessive-compulsive disorder. Neuron, 28:343–347

Gunaydin, LA, Kreitzer, AC (2016) Cortico-basal ganglia circuit function in psychiatric disease. Annual Review of Physiology, 78:327–350

Haber, SN (2016) Integrative networks across basal ganglia circuits. In Steiner, H and Tseng, K (Eds.): Handbook of basal ganglia structure and function, second edition. Elsevier. doi: http://dx.doi.org/10.1016/B978-0-12-802206-1.00027-1

Haber, SN (2003) The primate basal ganglia: parallel and integrative networks. Journal of Chemical Neuroanatomy, 26:317–330

Haroon, E, Fleischer, CC, Chen, X, Woolwine, BJ, Patel, T, Hu, XP, Miller, AH (2016) Conceptual convergence: increased inflammation is associated with increased basal ganglia glutamate in patients with major depressive disorder. Molecular Psychiatry, 21:1351–1357

Harrington, DL, Liu, D, Smith, MM, Mills, JA, Long, JD, Aylward, EH, Paulsen, JS, PREDICT-HD Investigators Coordinators of the Huntington Study Group (2014) Neuroanatomical correlates of cognitive functioning in prodromal Huntington disease. Brain and Behavior, 4(1):29–40

Harrington, DL, Rubinov, M, Durgerian, S, Mourany, L, Reece, C, Koenig, K, Bullmore, E, Long, JD, Paulsen, JS, Rao, SM (2015) Network topology and functional connectivity disturbances precede the onset of Huntington’s disease. Brain, 138:2332–2346

Hazrati, L-N, Parent, A (1992) The striatopallidal projection displays a high degree of anatomical specificity in the primate. Brain Research, 592:213–227

Hedreen, JC, DeLong, MR (1991) Organization of striatopallidal, striatonigral, and nigrostriatal projections in the macaque. J Comp Neurol, 304:569–595

Heemskerk, A-W, Roos, ACR (2011) Dysphagia in Huntington’s disease: a review. Dysphagia, 26:62–66

Heinsen, H, Rub, U, Bauer, M, Ulmar, G, Bethke, B, Schuler, M, Bocker, F, Eisenmenger, W, Gotz, M, Korr, H, Schmitz, C (1999) Nerve cell loss in the thalamic mediodorsal nucleus in Huntington’s disease. Acta Neuropathologica, 97:613–622

Hintiryan, H, Foster, NN, Bowman, I, Bay, M, Song, MY, Gou, L, Yamashita, S, Bienkowski, MS, Zingg, B, Zhu, M, Yang, XW, Shih, JC, Toga, AW, Dong, H-W (2016) The mouse cortico-striatal projectome. Nature Neuroscience, 19:1100–1114

Hoover, JE, Strick, PL (1993) Multiple output channels in the basal ganglia. Science, 259:819–821

Hunnicutt, BJ, Jongbloets, BC, Birdsong, WT, Gertz, KJ, Zhong, H, Mao, T (2016) A comprehensive excitatory input map of the striatum reveals novel functional organization. eLife, 5:e19103

Jankovic, J, Roos, RAC (2014) Chorea associated with Huntington’s disease: to treat or not to treat? Movement Disorders. 29(11):1414–1418

Kassubek, H, Juengling, FD, Ecker, D, Landwehrmeyer, GB (2005) Thalamic atrophy in Huntington’s disease co-varies with cognitive performance: a morphometric MRI analysis. Cerebral Cortex, 15:846–853

Kawaguchi, Y, Wilson, CJ, Emson, PC (1990) Projection subtypes of rat neostriatal matrix cells revealed by intracellular injection of biocytin. Journal of Neuroscience, 10:3421–3438

Kiferle, L, Mazzucchi, S, Unti, E, Pesaresi, I, Fabbri, S, Nicoletti, V, Volterrani, D, Cosottini, M, Bonuccelli, U, Ceravolo, R (2013) Parkinsonism and Related Disorders, 19:800–805

Kitano, H, Tanibuchi, I, Jinnai, K (1998) The distribution of neurons in the substantia nigra pars reticulata with input from the motor, premotor, and prefrontal areas of the cerebral cortex in monkeys. Brain Research, 784:228–238

Koob, GF, Volkow, ND (2016) Neurobiology of addiction: a neurocircuitry analysis. Lancet Psychiatry, 3(8):760–773

Kunzle, H (1978) An autoradiographic analysis of the efferent connections from premotor and adjacent prefrontal regions (areas 6 and 9) in *Macaca fascicularis*. Brain Behavior and Evolution, 15:185–234

Lancichinetti, A, Fortunato, S (2012) Consensus clustering in complex networks. Scientific Reports, 2:1–7

Levesque, M, Parent, A (2005) The striatofugal fiber system in primates: a reevaluation of its organization based on single-axon tracing studies. PNAS, 102(33):11888–11893

Levy, R, Dubois, B (2006) Apathy and the functional anatomy of the prefrontal cortex-basal ganglia circuits. Cerebral Cortex, 16:916–928

Li, Y, Gong, H, Yang, X, et al. (2017) TDat: an efficient platform for processing petabyte-scale whole-brain volumetric images. Frontiers in Neural Circuits, 11:51.

Lusk, NA, Buonomano, DV (2016) Utilizing the cortico-striatal projectome to advance the study of timing and time perception. Timing & Time Perception, 4(4):411–422, doi.org/10.1163/22134468-00002076

Mallet, N, Micklem, BR, Henny, P, Brown, MT, Williams, C, Bolam, JP, Nakamura, KC, Magill, PJ (2012) Dichotomous organization of the external globus pallidus. Neuron, 74(6):1075–1086

Mandelbaum, G, Taranda, J, Haynes, TM, Hochbaum, DR, Huang, KW, Hyun, M, Venkataraju, KU, Straub, C, Wang, W, Robertson, K, Osten, P, Sabatini, B (2019) Distinct cortico-thalamic-striatal circuits through the parafscicular nucleus. Neuron, 102(3):636–652

Mastro, KJ, Bouchard, RS, Holt, HAK, Gittis, AH (2014) Transgenic mouse lines subdivide external segment of the globus pallidus (GPe) neurons and reveal distinct GPe output pathways. Journal of Neuroscience, 34(6):2087–2099

Maurice, N, Deniau, J-M, Glowinski, J, Thierry, A-M (1999) Relationships between the prefrontal cortex and the basal ganglia in the rat: physiology of the cortico-nigral circuits. Journal of Neuroscience, 19(11):4674–4681

Middleton, FA, Strick, PL (2001) A revised neuroanatomy of frontal-subcortical circuits. In, Lichter, DG, Cummings, JL (Eds.) Frontal-subcortical circuits in psychiatric and neurological disorders. The Guilford Press: New York

Middleton, FA, Strick, PL (2002) Basal-ganglia ‘projections’ to the prefrontal cortex of the primate. Cerebral Cortex, 12:926–935

Naito, A, Kito, H (1994) The cortico-nigral projection in the rat: an anterograde tracing study with biotinylated dextran amine. Brain Research 637:317–322

Nambu, A, Takada, M, Inase, M, Tokuno, H (1996) Dual somatotopical representations in the primate subthalamic nucleus: evidence for ordered but reversed body-map transformations from the primary motor cortex and the supplementary motor area. Journal of Neuroscience, 16(8):2671–2683

Nambu, A, Tokuno, H, Hamada, I, Kita, H, Imanishi, M, Akazawa, T, Ikeuchi, Y, Hasegawa, N (2000) Excitatory cortical inputs to pallidal neurons via the subthalamic nucleus in the monkey. Journal of Neurophysiology, 84:289–300

Nambu, A, Tokuno, H, Takada, M (2002) Functional significance of the cortico-subthalamo-pallidal ‘hyperdirect’ pathway. Neuroscience Research, 43:111–117

Nambu, A, Yoshida, S, Jinnai, K (1988) Projection on the motor cortex of thalamic neurons with pallidal input in the monkey. Exp Br Research, 71:658–662

Nambu, A, Yoshida, S, Jinnai, K (1991) Movement-related activity of thalamic neurons with input from the globus pallidus and projection to the motor cotex in the monkey. Exp Br Res, 84:279–284

Nauta, WJH, Mehler, WR (1961) Some efferent connections of the lentiform nucleus in monkey and cat. The Anatomical Record, 139:260

Navone, F, Jahn, R, Di Gioia, G, Stukenbrok, H, Greengard, P, De Camilli, P (1986) Protein 38: an integral membrane protein specific for small vesicles of neurons and neuroendocrine cells. J Cell Biol, 103(6):2511–2527

Nestler, EJ, Barrot, M, DiLeone, RJ, Eisch, AJ, Gold, SJ, Monteggia, LM (2002) Neurobiology of depression. Neuron, 34:13–25

Ni, H, Tan, C, Feng, Z, Chen, S, Zhang, Z, Li, W, Guan, Y, Gong, H, Luo, Q, Li, Anan (2020) A robust image registration interface for large volume brain atlas. Scientific Reports, 10:2139

Oh, SW, Harris, JA, Ng, L, Winslow, B, Cain, N, Mihalas, S … Jones, AR, Zeng, H (2014) A mesoscale connectome of the mouse brain. Nature, 508:207–214

O’Hare, J, Calakos, N, Yin, HH (2018) Recent insights into corticostriatal circuit mechanisms underlying habits. Current Opinion in Behavioral Sciences, 20:40–46

Oyanagi, K, Takeda, S, Takahashi, H, Ohama, E, Ikuta, F (1989) A quantitative investigation of the substantia nigra in Huntington’s disease. Annals of Neurology, 26:13–19

Parent, A, Hazrati, L-N (1994) Multiple striatal representation in primate substantia nigra. Journal of Comparative Neurology, 344:305–320

Parent, A, Hazrati, L-N (1995) Functional anatomy of the basal ganglia. I. The cortico-basal ganglia-thalamo-cortical loop. Brain Research Reviews, 20:91–127

Park, Y-G, Sohn, CH, Chen, R, McCue, M, Yun, DH, Drummond, GT, Ku, T, Evans, NB, Oak, HC, Trieu, W, Choi, H, Jin, X, Lilascharoen, V, Wang, J, Truttman, MC, Qi, HW, Ploegh, HL, Golub, TR, Chen, S-C, Frosch, MP, Kulik, HJ, Lim, BK, Chung, K (2019) Protection of tissue physiochemical properties using polyfunctional crosslinkers. Nature Biotechnology, 37(1):73–83 doi: 10.1083/nbt.4281

Paulsen, JS, Miller, AC, Hayes, T, Shaw, E (2017) Cognitive and behavioral changes in Huntington disease before diagnosis. Handbook of Clinical Neurology, 144:69–91

Petreanu, L, Huber, D, Sobczyk, A, Svoboda, K (2007) Channelrhodopsin-2-assisted circuit mapping of long-range callosal projections. Nature Neuroscience, 10(5):663–668

Pievani, M, de Haan, W, Wu, T, Seeley, WW, Frisoni, GB (2011) Functional network disruption in the degenerative dementias. Lancet Neurology, 10:829–843

Rinvik, E, Walberg, F (1969) Is there a cortico-nigral tract? A comment based on experimental electron microscopic observations in the cat. Brain Research, 14:742–744

Robledo, P, Feger, J (1990) Excitatory influence of rat subthalamic nucleus to substantia nigra pars reticulata and the pallidal complex: electrophysiological data. Brain Research, 518(1-2):47–54

Rosas, HD, Koroshetz, WJ, Chen, YI, Skeuse, C, Vangel, M, Cudkowicz, ME, Caplan, K, Marek, K, Seidman, LJ, Makris, N, Jenkins, BG, Goldstein, JM (2003) Evidence for more widespread cerebral pathology in early HD. Neurology, 60:1615–1620

Rosas, HD, Tuch, DS, Hevelone, ND, Zaleta, AK, Vangel, M, Hersch, SM, Salat, DH (2006) Diffusion tensor imaging in presymptomatic and early Huntington’s disease: selective white matter pathology and its relationship to clinical measures. Movement Disorders, 21(9):1317–1325

Rouiller, EM, Liang, F, Babalian, A, Moret, V, Wiesendanger, M (1994) Cerebellothalamocortical and pallidothalamocortical projections to the primary and supplementary motor cortical areas: a multiple tracing study in macaque monkeys. Journal of Comparative Neurology, 345:185–213

Sato, F, Lavallee, P, Levesque, M, Parent, A (2000) Single-axon tracing study of neurons of the external segment of the globus pallidus in primate. Journal of Comparative Neurology, 417:17–31

Saunders, A, Oldenburg, IA, Berezovskii, VK, Johnson, CA, Kingery, ND, Elliott, HL, Xie, T, Gerfen, CR, Sabatini, BL (2015) A direct GABAergic output from the basal ganglia to frontal cortex. Nature, 521:85–89

Schultz, W (1986) Activity of pars reticulata neurons of monkey substantia nigra in relation to motor, sensory, and complex events. J Neurophysiology, 55(4):660–677

Seeley, WW, Crawford, RK, Zhou, J, Miller, BL, Greicius, MD (2009) Neurodegenerative diseases target large-scale human brain networks. Neuron, 62:42–52

Simmons, DV, Higgs, MH, Lebby, S, Wilson, CJ (2020) Indirect pathway control of firing rate and pattern in the substantia nigra pars reticulata. Journal of Neurophysiology, 123:800–814

Smith, Y, Bolam, JP (1989) Neurons of the substantia nigra reticulata receive a dense GABA-containing input from the globus pallidus in the rat. Brain Research, 493:160–167

Smith, Y, Bolam, JP (1991) Convergence of synaptic inputs from the striatum and the globus pallidus onto identified nigrocollicular cells in the rat: a double anterograde labeling study. Neuroscience, 44(1):45–73

Smith, KS, Graybiel, AM (2014) Investigating habits: strategies, technologies and models. Frontiers in Behavioral Neuroscience, 8(39):1–17 doi: 10.3389/fnbeh.2014.00039

Sun, P, Jin, S, Tao, S, Wang, J, Li, A, Wu, Y, Kuang, J, Liu, Y, Wang, L, … Zhang, Y-H, Xu, F (2019) Recombinase system-dependent copackaging strategy for highly efficient neurocircuit tracing. bioRxiv doi: https://doi.org/10.1101/705772

Swanson, LW (2000) Cerebral hemisphere regulation of motivated behavior. Brain Research, 886:113–164

Szabo, J (1962) Topical distribution of the striatal efferents in the monkey. Experimental Neurology, 5:21–36

Tabrizi, SJ, Scahill, RI, Durr, A, Roos, RAC, Leavitt, BR, Jones, R, Landwehrmeyer, GB, Fox, NC, Johnson, H, Hicks, SL, Kennard, C, Craufurd, D, Frost, C, Langbehn, DR, Reilmann, R, Stout, JC, The TRACK-HD investigators (2011) Biological and clinical changes in premanifest and early stage Huntington’s disease in the TRACK-HD study: the 12-month longitudinal analysis. Lancet Neurology, 10(1):31–42

Tanaka, SC, Balleine, BW, O’Doherty, JP (2008) Calculating consequences: brain systems that encode the causal effects of actions. Journal of Neuroscience, 28(26):6750–6775

Tanaka, YH, Tanaka, YR, Kondo, M, Terada, S-I, Kawaguchi, Y, Matsuzaki, M (2018) Thalamocortical axonal activity in motor cortex exhibits layer-specific dynamics during motor learning. Neuron, 100:244–258

Tecuapetla, F, Jin, X, Lima, SQ, Costa, RM (2016) Complementary contributions of striatal projection pathways to action initiation and execution. Cell, 166:703–715

Thompson, RH, Swanson, LW (2010) Hypothesis-driven structural connectivity analysis supports network over hierarchical model of brain architecture. PNAS, 107(34):15235–15239

Tricomi, E, Balleine, BW, O’Doherty, JP (2009) A specific role for posterior dorsolateral striatum in human habit learning. European Journal of Neuroscience, 29:2225–2232

Vaghi, MM, Vertes, PE, Kitzbichler, MG, Apergis-Schoute, AM, van der Flier, FE, Fineberg, NA, Sule, A, Zaman, R, Voon, V, Kundu, P, Bullmore, ET, Robbins, TW (2017) Specific frontostriatal circuits for impaired cognitive flexibility and goal-directed planning in obsessive-compulsive disorder: evidence from resting state functional connectivity. Biological Psychiatry, 81:708–717

Vonsattel, JPG, DiFiglia, M (1998) Huntington disease. Journal of Neuropathology and Experimental Neurology, 57(5):369–384

Wall, NR, De La Parra, M, Callaway, EM, Kreitzer, AC (2013) Differential innervation of direct- and indirect-pathway striatal projection neurons. Neuron, 79:347–360

Wallace, ML, Saunders, A, Huang, KW, Philson, AC, Goldman, M, Macosko, EZ, McCarroll, SA, Sabatini, B (2017) Genetically distinct parallel pathways in the entopeduncular nucleus for limbic and sensorimotor output of the basal ganglia. Neuron, 94:138–152

Whiteside, SP, Port, JD, Abramowitz, JS (2004) A meta-analysis of functional neuroimaging in obsessive-compulsive disorder. Psychiatry Research: Neuroimaging, 132:69–79

Wiedenmann, B, Franke, WW (1985) Identification and localization of synaptophysin, an integral membrane glycoprotein of M_r_ 38,000 characteristic of presynaptic vesicles. Cell, 41(3):P1017–1028

Wu, Y, Richard, S, Parent, A (2000) The organization of the striatal output system: a single-cell juxtacellular labeling study in the rat. Neuroscience Research, 38:49–62

Yin, HH, Knowlton, BJ, Balleine, BW (2004) Lesions of dorsolateral striatum preserve outcome expectancy but disrupt habit formation in instrumental learning. European Journal of Neuroscience, 19:181–189

Yin, HH, Ostlund, SB, Knowlton, BJ, Balleine, BW (2005) The role of the dorsomedial striatum in instrumental conditioning. European Journal of Neuroscience, 22:513–523

Zhou, H, Li, S, Li, A, Xiong, F, Li, N, Han, J…Quan, T, Zeng, S (2018) Dense reconstruction of brain-wide neuronal population close to the ground truth. bioRxiv, doi: https://doi.org/10.1101/223834

Zingg, B, Chou, X, Zhang, Z, Mesik, L, Liang, F, Tao, HW, Zhang, LI (2017) AAV-mediated anterograde transsynaptic tagging: mapping corticocollicular input-defined neural pathways for defense behavior. Neuron, 93(1):33–47

Zingg, B, Hintiryan, H, Gou, L, Song, MY, Bay, M, Bienkowski, MS, Foster, NN, Yamashita, S, Bowman, I, Toga, AW, Dong, H-W (2014) Neural networks of the mouse neocortex. Cell, 156:1096–1111

Zingg, B, Peng, B, Huang, JJ, Tao, HW, Zhang, LI (2020) Synaptic specificity and application of anterograde transsynaptic AAV for probing neural circuitry. Journal of Neuroscience, 40(16):3250–3267

